# Catch composition and life history characteristics of sharks and rays (Elasmobranchii) landed in the Andaman and Nicobar Islands, India

**DOI:** 10.1101/2020.03.17.995217

**Authors:** Zoya Tyabji, Tanmay Wagh, Vardhan Patankar, Rima W. Jabado, Dipani Sutaria

## Abstract

The scientific literature on the diversity and biological characteristics of sharks and rays from the Andaman and Nicobar Archipelago fishing grounds is scarce and compromised by species misidentifications. We carried out systematic fish landing surveys in South Andamans from January 2017 to May 2018, a comprehensive and cost-effective way to fill this data gap. We sampled 5,742 individuals representing 57 shark and ray species. Of the 36 species of sharks and 21 species of rays landed, six species of sharks - *Loxodon macrorhinus, Carcharhinus amblyrhynchos, Sphyrna lewini, Carcharhinus albimarginatus*, *Carcharhinus brevipinna,* and *Paragaelus randalli* dominated landings and comprised 83.35 % of shark landings, while three species of rays were most abundant – *Pateobatis jenkinsii*, *Himantura leoparda* and *H. tutul*, and comprised 48.82 % of ray landings. We report size extensions for seven shark species as well as three previously unreported ray species, increasing the known diversity for the islands and for India. For sharks, mature individuals of small-bodied species (63.48 % males of total landings of species less than 1.5 m total length) and immature individuals of larger species (84.79 % males of total landings of species larger than 1.5 m total length) were mostly landed; whereas for rays, mature individuals were predominantly landed (80.71 % males of total landings) likely reflecting differences in fishing patterns as well as habitat preferences and life history stages across species. Further, juvenile sharks and gravid females were landed in large quantities which might be unsustainable in the long-term. Landings were female-biased in *C. amblyrhynchos, S. lewini* and *P. jenkinsii,* and male-biased in *L. macrorhinus* and *H. leoparda*, indicating either spatio-temporal or gear specific sexual segregation in these species. Understanding these nuances - the composition and biology of sharks and rays landed in different fisheries seasonally will inform future conservation and fishery management measures for these species in the Andaman and Nicobar Islands.

## INTRODUCTION

Elasmobranchs (sharks and rays) are recognized as one of the marine taxa with the highest extinction risk and need for urgent conservation measures [1]. Despite considerable inter and intra-specific life history variation [2, 3], most species have relatively low productivity making them highly susceptible to anthropogenic and natural stressors [4]. Populations of many species have drastically declined globally due to overfishing and habitat degradation raising concerns about their long-term survival [1].

In the past few decades, India has consistently been one of the top three shark and ray harvesters in the world [5, 6]. Here, sharks and rays are primarily caught as bycatch [7–11] in a large fishing fleet of 238,772 registered commercial and artisanal fishing crafts [12]. However, a few targeted shark fisheries that formed in the 1980’s remain including in the Andaman and Nicobar Islands [13, 14]. Anecdotal information from interviews with fishers on these islands indicate that shark and ray populations have declined [15] but there have been few systematic surveys of landings carried out to assess the current situation. This limited information on species and stocks may have detrimental effects not only on the ecology of these animals but also on the sustainability of these fisheries and the food security they provide as well as the socio-economic dependence of fisher communities [16, 17].

Over the years, with growing reports of declining populations of sharks and rays, the Government of India has implemented several conservation policies. In 2001, ten species of sharks and rays, including the Whale shark *Rhincodon typus,* Knifetooth sawfish *Anoxypristis cuspidata*, Pondicherry shark *Carcharhinus hemiodon,* Gangetic shark *Glyphis gangeticus,* Speartooth shark *G. glyphis,* Ganges stingray *Himantura fluviatilis,* Freshwater sawfish *Pristis microdon* (*= P. pristis*), Green sawfish *P. zijsron*, Giant guitarfish *Rhynchobatus djiddensis*, and Porcupine ray *Urogymnus asperrimus* were listed under Schedule I of the Indian Wildlife (Protection) Act, 1972 (WLPA). In 2009, the Andaman and Nicobar Islands Fisheries Regulation declared a 45-day closed season for shark fishing from April 15^th^ to May 31^st^ around the islands through the prohibition of the use of shark and tuna pelagic longlines and trawl nets. In 2013, the Ministry of Environment, Forest and Climate Change (MoEF&CC) declared a ‘Fin-attached Policy’ where sharks have to be landed whole, with their fins naturally attached to their bodies. In 2015, India’s Ministry of Commerce and Industry issued a notification prohibiting the export of all shark fins. While these legislations, if properly implemented, are a positive step for shark conservation in India, there appears to be an agenda mismatch between the MoEF&CC and the Ministry of Animal Husbandry, Dairying and Fisheries, with the latter having recently developed a strategy to expand fisheries and increase yield. This includes developing new schemes and projects to harness fishing potential and create employment opportunities, by issuing additional fishing licenses, building infrastructure such as cold storage centers, blast freezers and ice plants, and increasing introduction of deep-sea, motorized and mechanized boats [18].

In order to develop best management practices, basic life-history information such as age, growth, and maturity is required to form the basis of population assessments. However, in many developing countries, landings remain unmonitored and unregulated with little species-specific data collected, which hampers population assessments, and does not provide indication of the status of a population [17]. Additionally, since different species can show variances in biological traits across geographical regions, such as size at birth, size at maturity, maximum size, litter size, and breeding cycle [19–21], it is important to undertake region-specific studies so they can inform local management strategies.

Most past literature on sharks and rays from the Andaman and Nicobar Islands has been limited to species identification and taxonomy [22–27]. A large knowledge gap exists in our understanding of the catch composition of commercial species landed, their population trends, biological characteristics across seasons. Here, we aim to address this gap by assessing sharks and rays landed in the Andaman and Nicobar Islands and exploring 1) the species composition including relative abundance across seasons, 2) the biological information, including size frequency, sex ratio, maturity and length-weight relationships, and 3) the characteristics of fishing gears and grounds where sharks and rays were reportedly fished.

## MATERIALS AND METHODS

### Study area

The Andaman and Nicobar Islands (6°N–14°N and 92°E–94°E) are located in the Bay of Bengal and constitute 29.7 % of the total Economic Exclusive Zone (EEZ) area of India (Fig 1), covering a coastline of 1,962 km (contributing to 26.10 % of India’s coastline) and a continental shelf area of approximately 35,000 km^2^ [18]. The islands experience heavy monsoon from end of May to September when the south-west monsoon sets in as well as intermittent or light to heavy rainfall when the north-east monsoon sets in November. For the duration of our study, we characterized landings according to the following seasons: north-east monsoon (NE) (October–January), dry season (DS) (February–May), and south-west monsoon (SW) (June–September).

**Fig 1.**
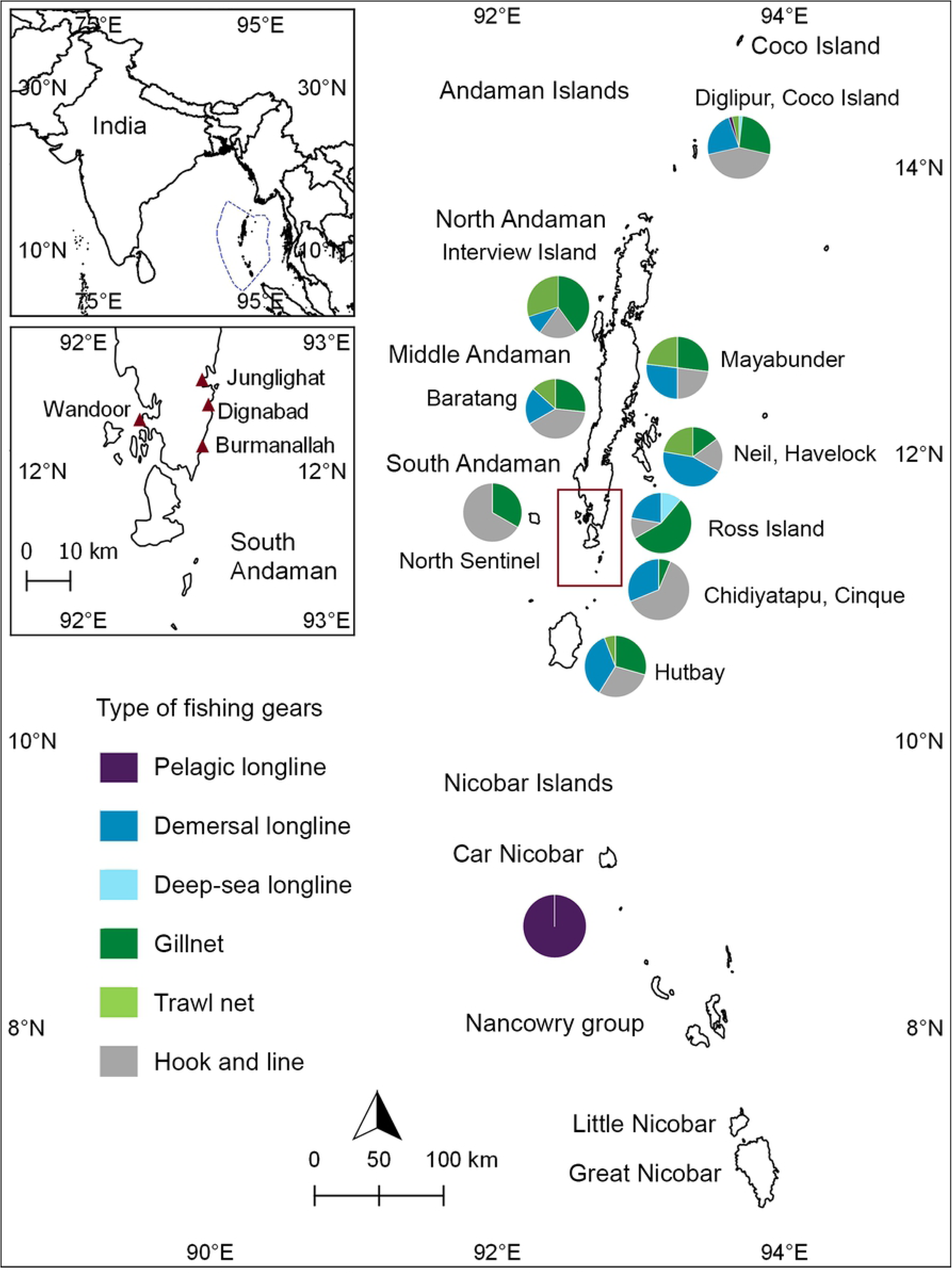
Map of the sampling sites and fishing grounds of the Andaman and Nicobar Islands, India. Top left: Map of India with the Exclusive Economic Zone boundaries of Andaman and Nicobar Islands demarcated in blue; Bottom left: Map of South Andaman with red triangles indicating sampled fish landing centers; Right: Map of the Andaman and Nicobar Islands showing fishing gear utilization across fishing grounds around the Islands, South Andaman is demarcated by the red inset.

A total of 2,784 fishing vessels are currently active with 7,034 licensed fishers [18]. Here, sharks and rays are targeted using pelagic and deep-sea longlines; and are caught as bycatch in demersal longlines, trawl nets, gillnets, and hook and line. Pelagic longliners and trawlers are permitted to exclusively fish beyond six nautical miles and up to 12 nautical miles from the coast. Demersal longliners as well as hook and line and gillnet fisheries operate near the coast and shallow seamounts. Fishers from the Andaman Islands fish across the waters of the Andaman and Nicobar Islands while the communities on the Nicobar Islands, due to their seclusion, only fish for subsistence or fishing for sale in local markets [27, 28].

Exploratory visits to landing sites in 2016 across South Andaman Islands revealed that the majority of sharks and rays fished throughout the Andaman and Nicobar Islands are landed at Junglighat (Fig 1). Junglighat, located in Port Blair, the main city of the Andaman Islands, is the largest fish landing center of the islands with proximity to storage centers and export facilities (Fig 1). We therefore focused our sampling at this location. However, opportunistic surveys were also undertaken at the fish landing sites of Burmanallah, Wandoor, and Dignabad (Fig 1) when fishers or informants reported landings of sharks and rays to the survey team.

### Sampling effort

Systematic surveys were undertaken from January 2017 to May 2018 for sharks and from October 2017 to May 2018 for rays. Junglighat was visited every alternate day or when weather permitted from 0600 to 1000 hrs, whereas the remaining site visits were dependent on reports by the informants. As the pelagic longliners from Junglighat directly offload and transport their landings to the processing and storage units, sampling of landings from these vessels was conducted at these units between 1000 to 1400 hrs.

Sharks and rays landed were identified to the species level using the available literature and photo-documented [29–32]. Rays were often landed with their tails cut, in piles, and, in a few cases, when landings were large, accurate pictures and/or measurements were not possible. Therefore, species which were difficult to differentiate morphologically, such as *Neotrygon* sp. and *Pastinachus* sp. were grouped at the genus level, and have therefore been excluded from the analysis of the full data set.

For sharks, guitarfishes, and wedgefishes, the total length (TL, a straight line from the tip of the snout to the tip of the tail, with tail flexed down to midline) was measured, whereas for rays, the disc width (DW, a straight line at the widest region of the disc) was measured to the nearest millimeter [30, 31].

Sex was determined by the presence of claspers indicative of males or the absence of claspers indicative of females. For males, the degree of calcification and length of claspers determined the maturity levels. This was categorized from 1 to 3 where (1) refers to immature individuals whose claspers were non-calcified and pliable, and whose length was less than the pelvic fins, (2) refers to maturing individuals whose claspers were partially calcified and semi-pliable, and whose length was longer than the pelvic fins, and (3) refers to fully mature individuals whose claspers were fully calcified and non-pliable.

Gravid females were recorded by the presence of emerging embryos or if these could be clearly observed by pressing the stomach. Whenever possible, gravid females were dissected to record the sex and size of embryos. Young-of-the-year (YOY) individuals were identified by the presence of open umbilical scars which usually close after the first few months of life.

Weights were recorded to the nearest gram using a hand-held digital weighing balance for smaller individuals or whenever possible, when weights were provided by the fishers using a circular weighing balance for larger individuals (> 50 kg).

For each boat that landings were sampled from, we approached fishers for information on the fishing gears used to catch the sharks and rays, and the fishing grounds.

### Data analysis

A species accumulation curve over time, trends in mean abundance of landings across months, and patterns in species, sex, and sizes caught across various gears were produced in Microsoft Excel 2017. Tentative fishing grounds, including usage of fishing gears, were mapped on QGIS based on locations provided during discussions with fishers.

The hypothesis of equal sex ratios for species where ≥ 50 individuals were sampled, was tested using Chi-square where significance was considered at p < 0.05 [33]. The hypothesis of shark TL getting equally caught across different fishing gears was tested using one-way ANOVA where significance was considered at p < 0.05 [33].

For sharks, species where > 150 individuals were sampled, size-class frequency distributions by sex and seasons were plotted while the size at 50 % maturity (TL_50_) for males was calculated. This was done by fitting the following logistic function to the proportion of mature individuals in 10 or 20 cm size categories: P = 1 / (1 + exp (−r (LT_mid_ − LT_50_))), where P is the proportion of mature fish in each length class, LT_mid_ is the midpoint of the length class, LT_50_ is the mean size at sexual maturity, and r is a constant that increases in value with the steepness of the maturation schedule.

Finally, length-weight relationships were determined using regression analysis. The equation W=aL^b^ was converted into a linear form In (W) = In (a) + b In (L), where W is the weight, L is the length, ln(a) is the intercept and (b) the slope or regression coefficient. Gravid females were excluded from this analysis.

## RESULTS

Sampling was conducted on 216 days with landings recorded from 567 boats and a total of 5,742 sharks and rays representing 57 species. Of these, 4,632 individuals represented 36 shark species from 18 genera and 11 families while 1,110 individuals represented 21 ray species, 14 genera and eight families.

The species accumulation curve reached a threshold for sharks but not for rays (Fig 2). A species list and summary of biological data for all specimens is provided in Table 1.

**Fig 2.**
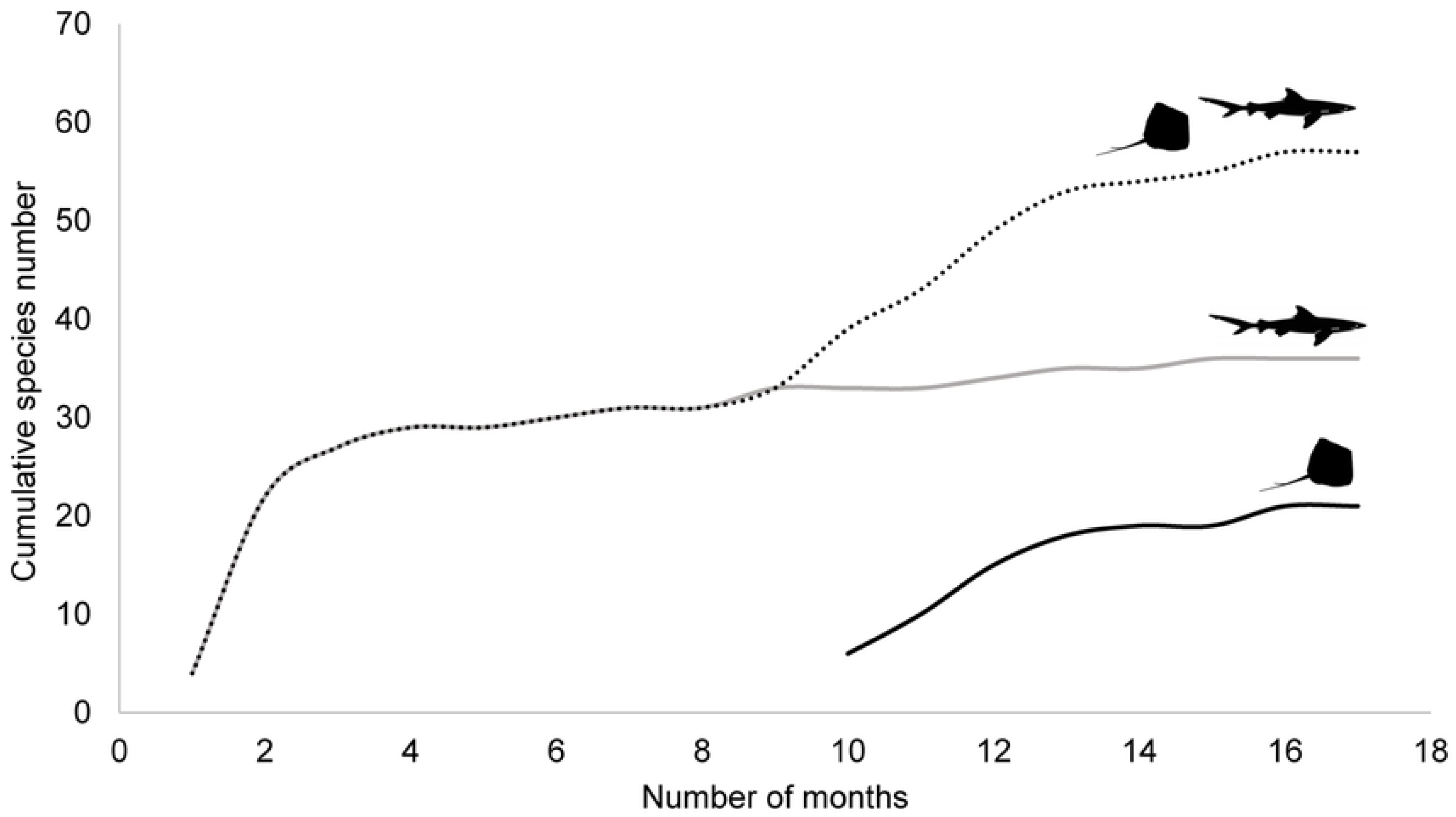
Species accumulation curve of sharks and rays landed. The curve shows the number of cumulative species across 17 months of sampling from January 2017 to May 2018. The dotted line represents both shark and ray species, the grey line represents shark species and the black line represents ray species.

**Table 1.**
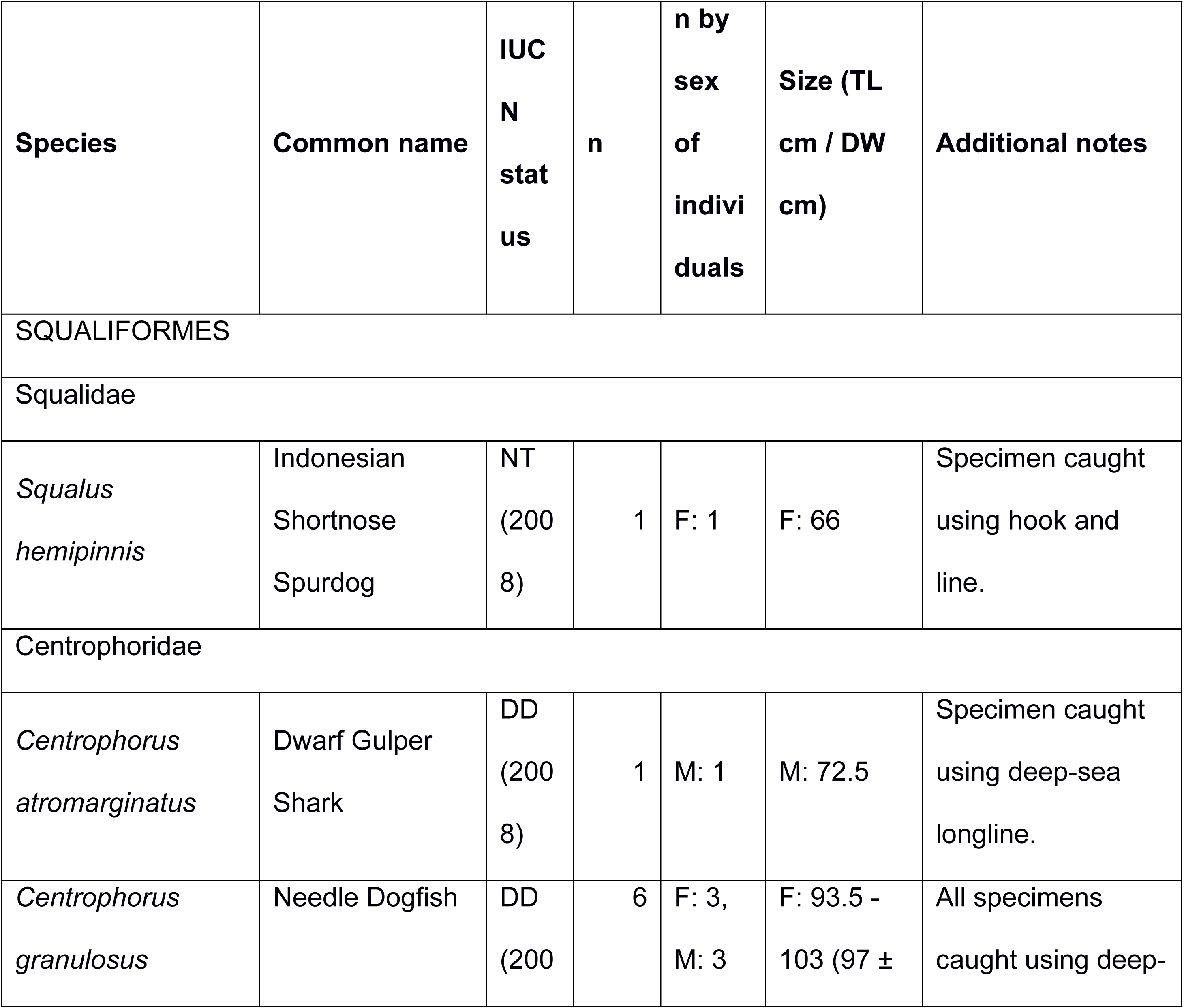
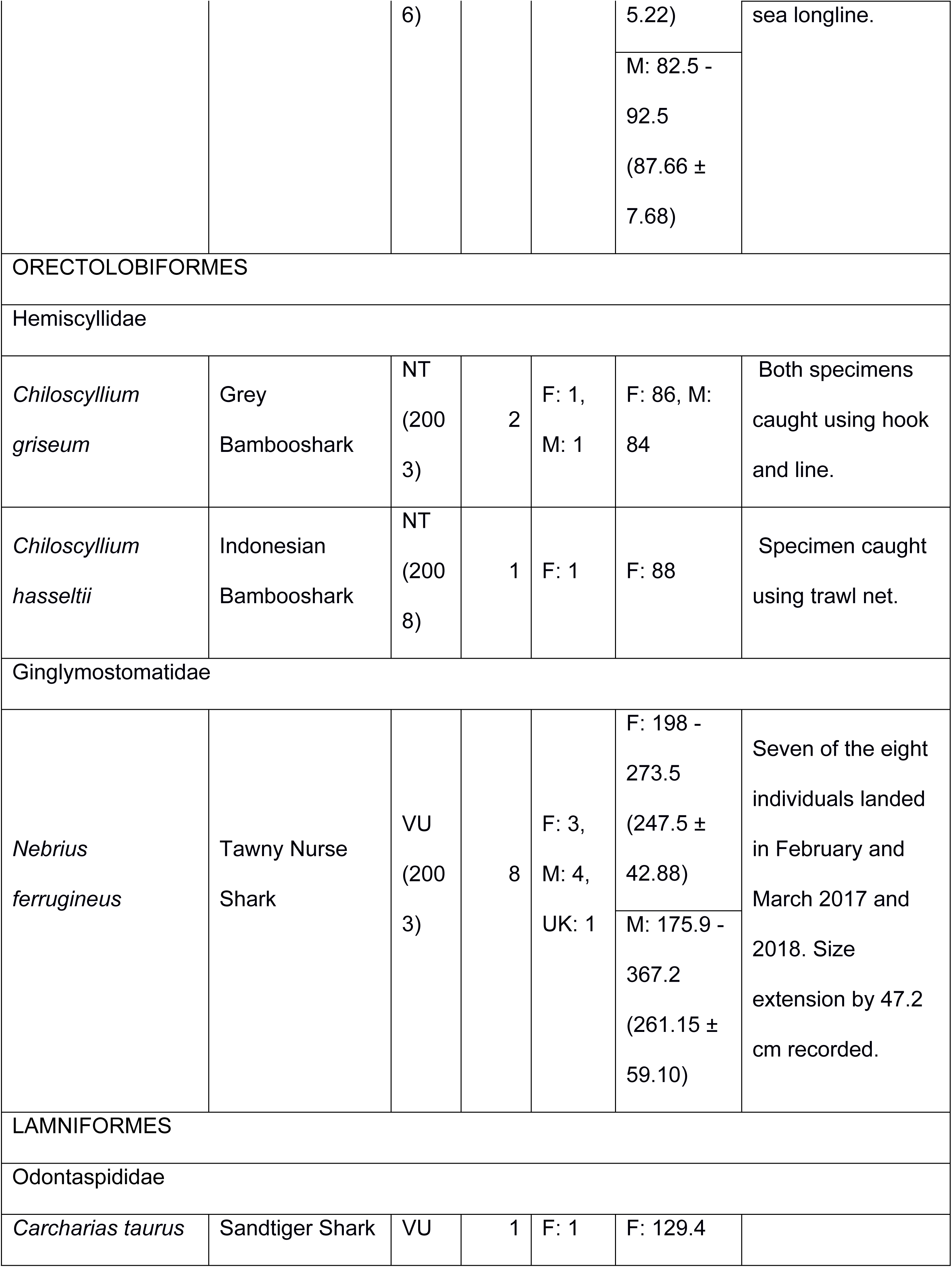
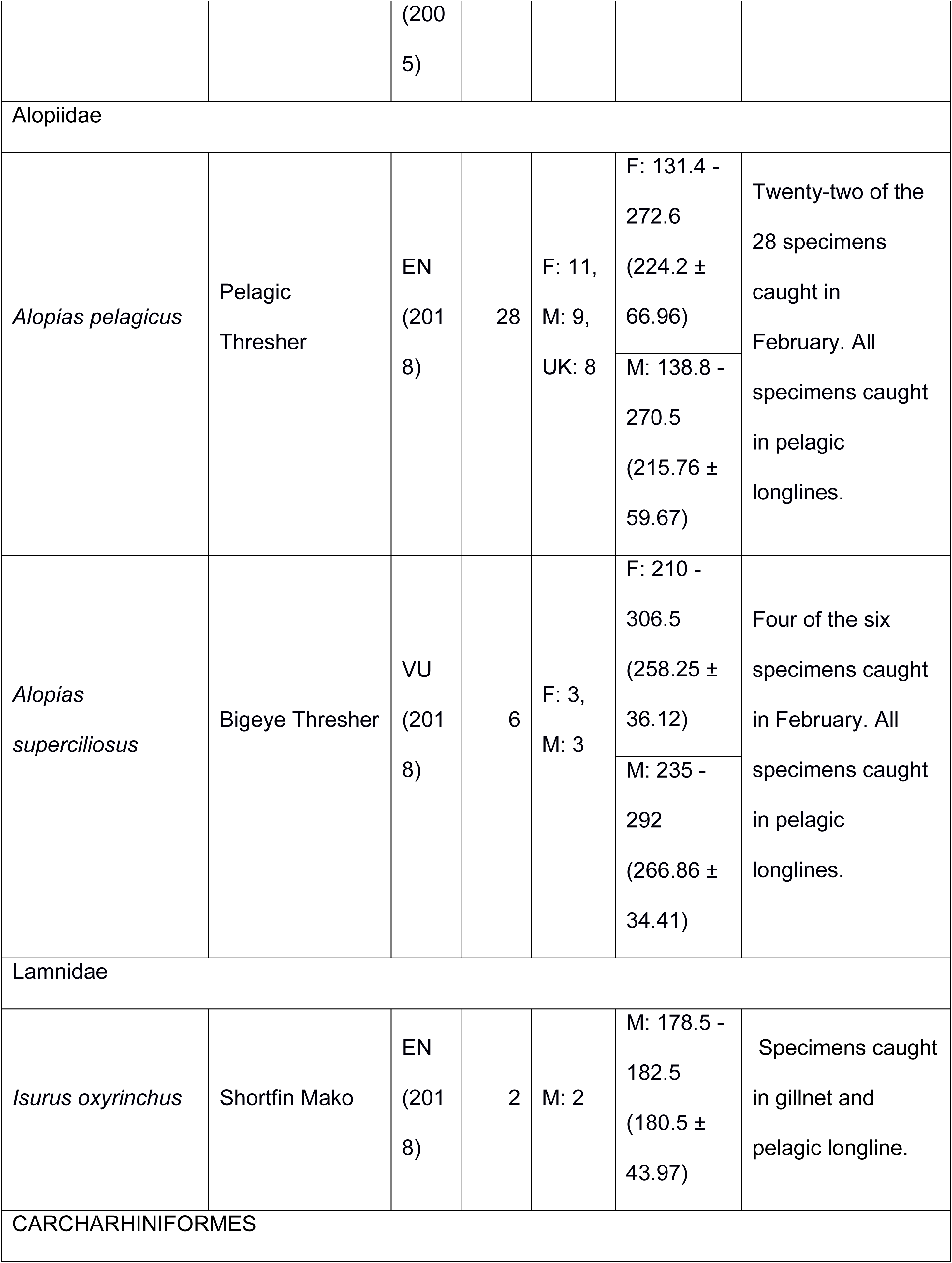
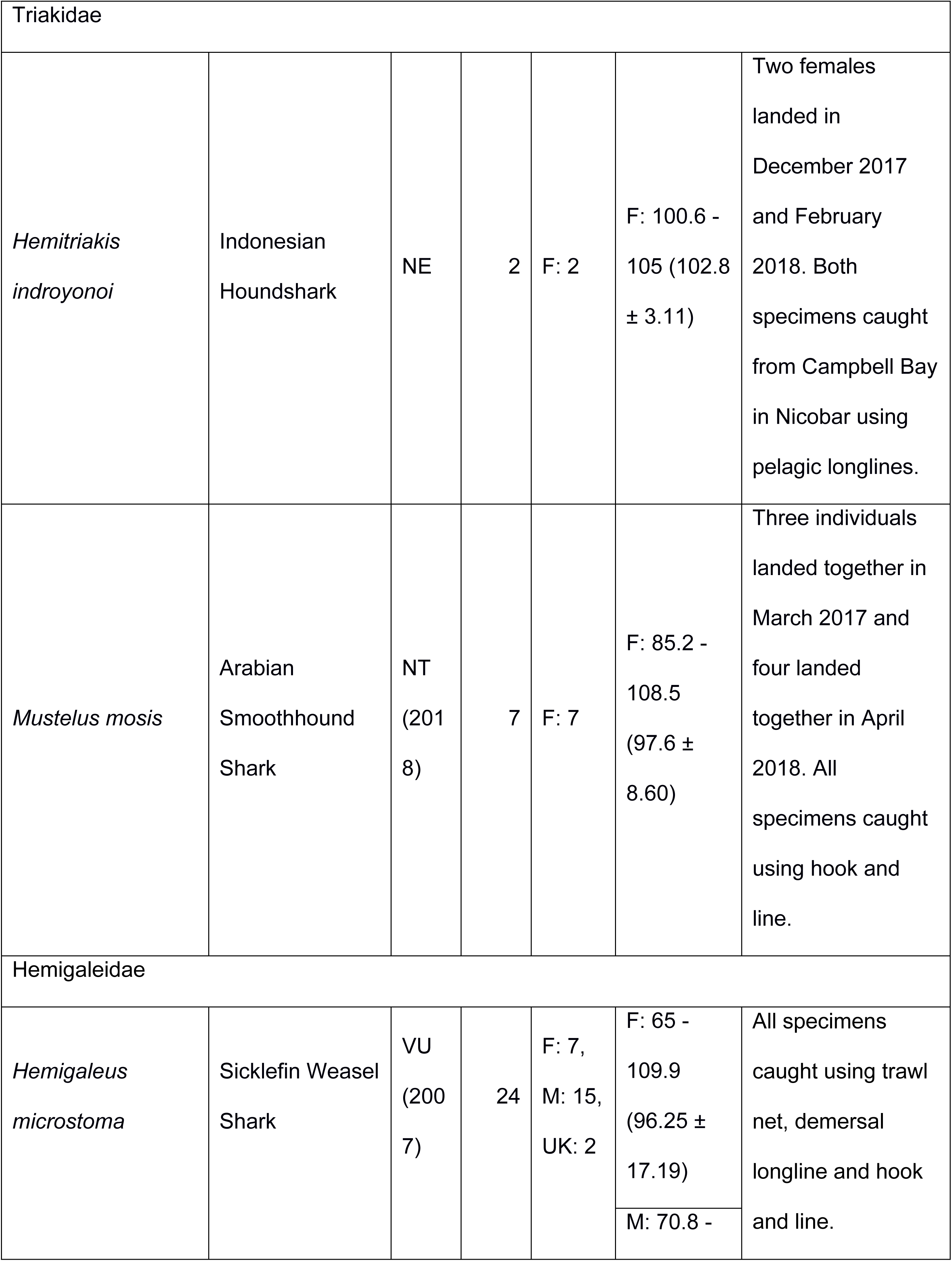
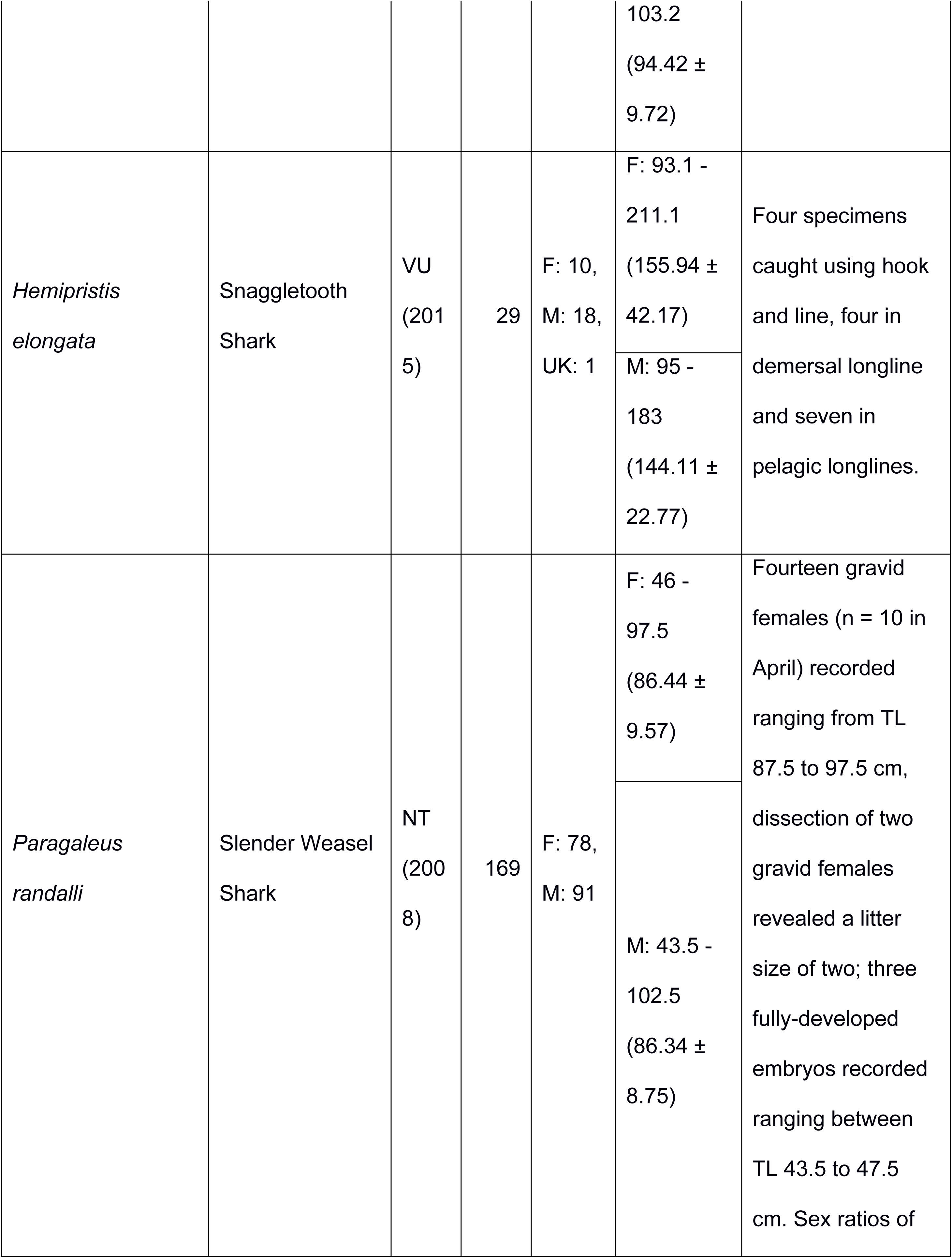
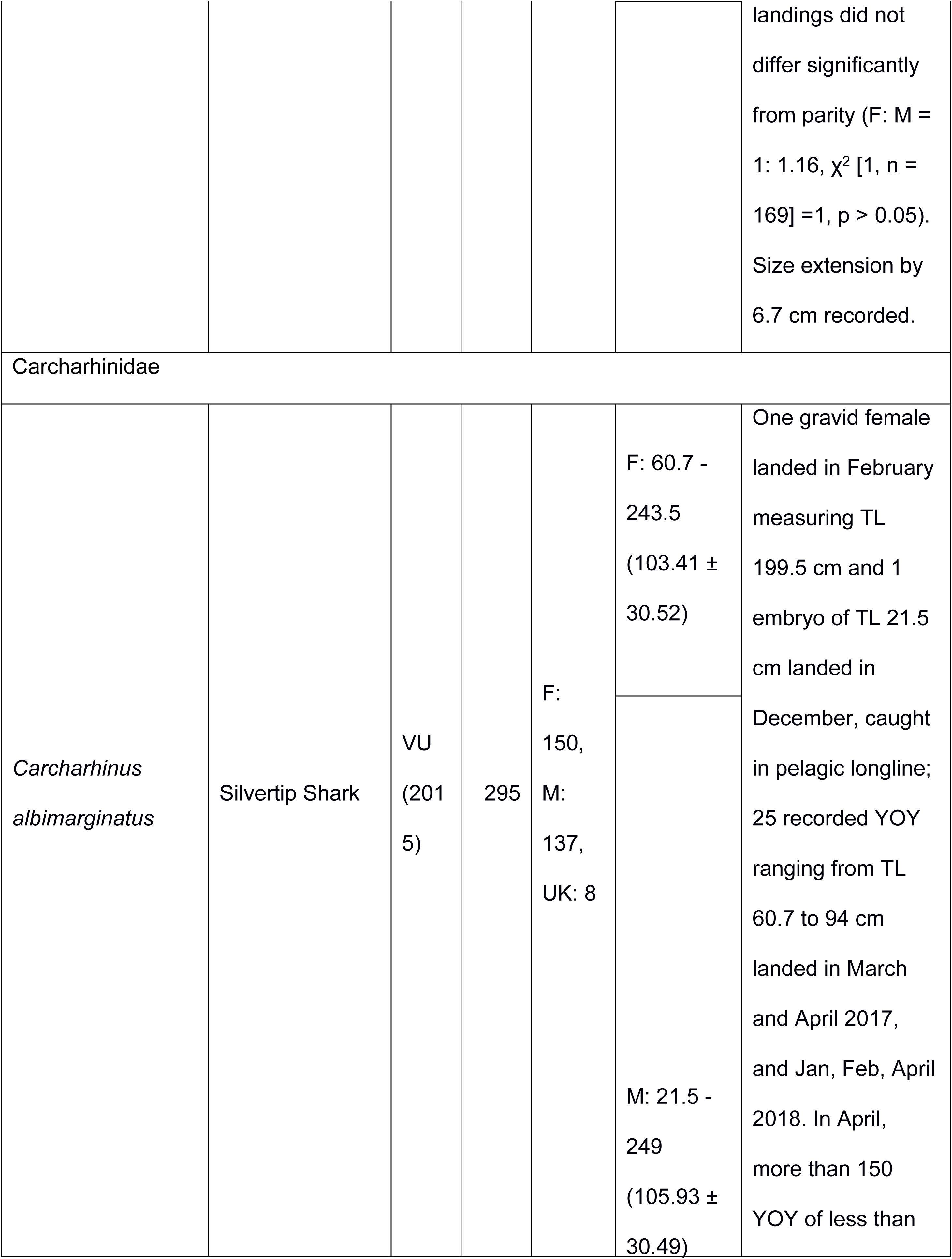
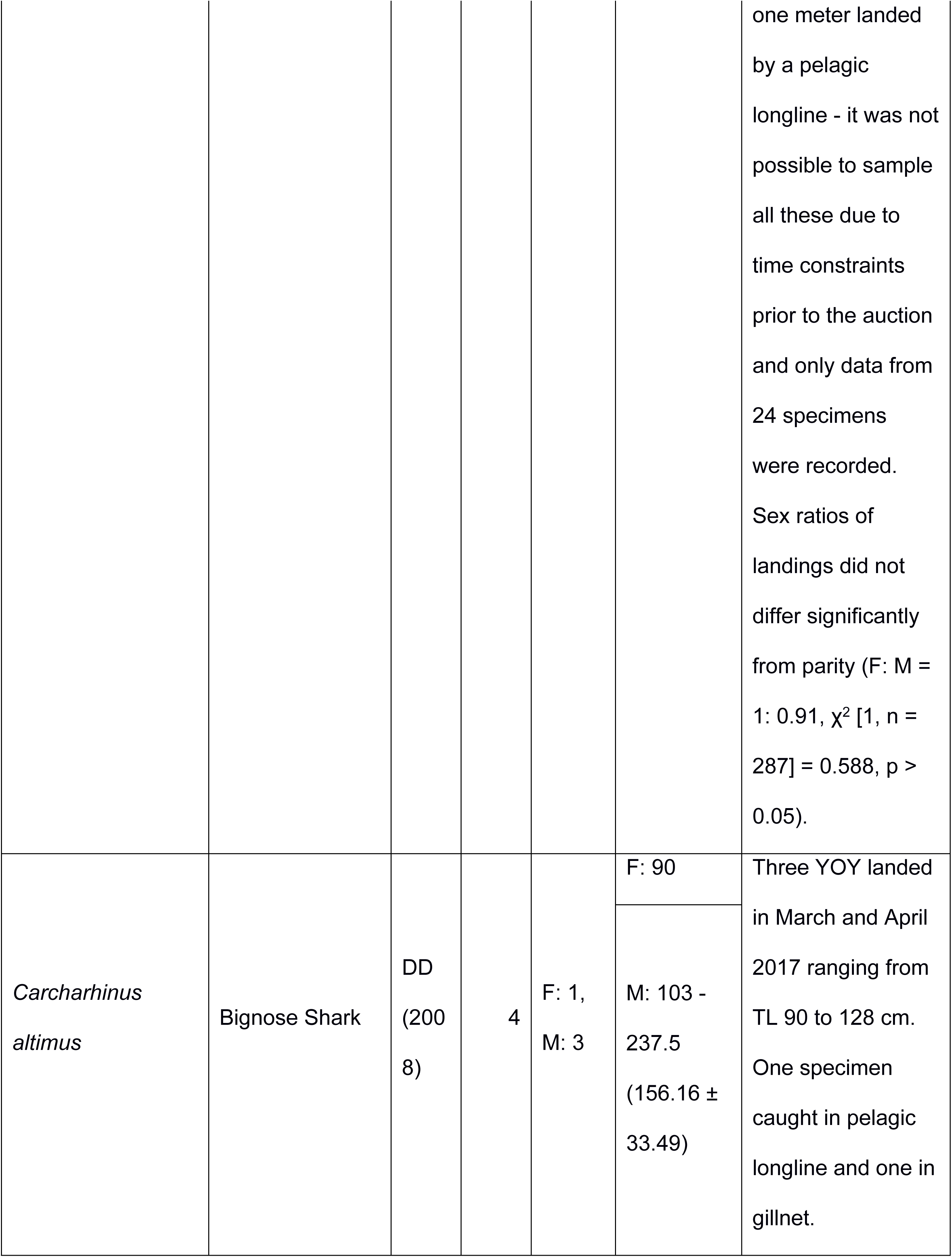
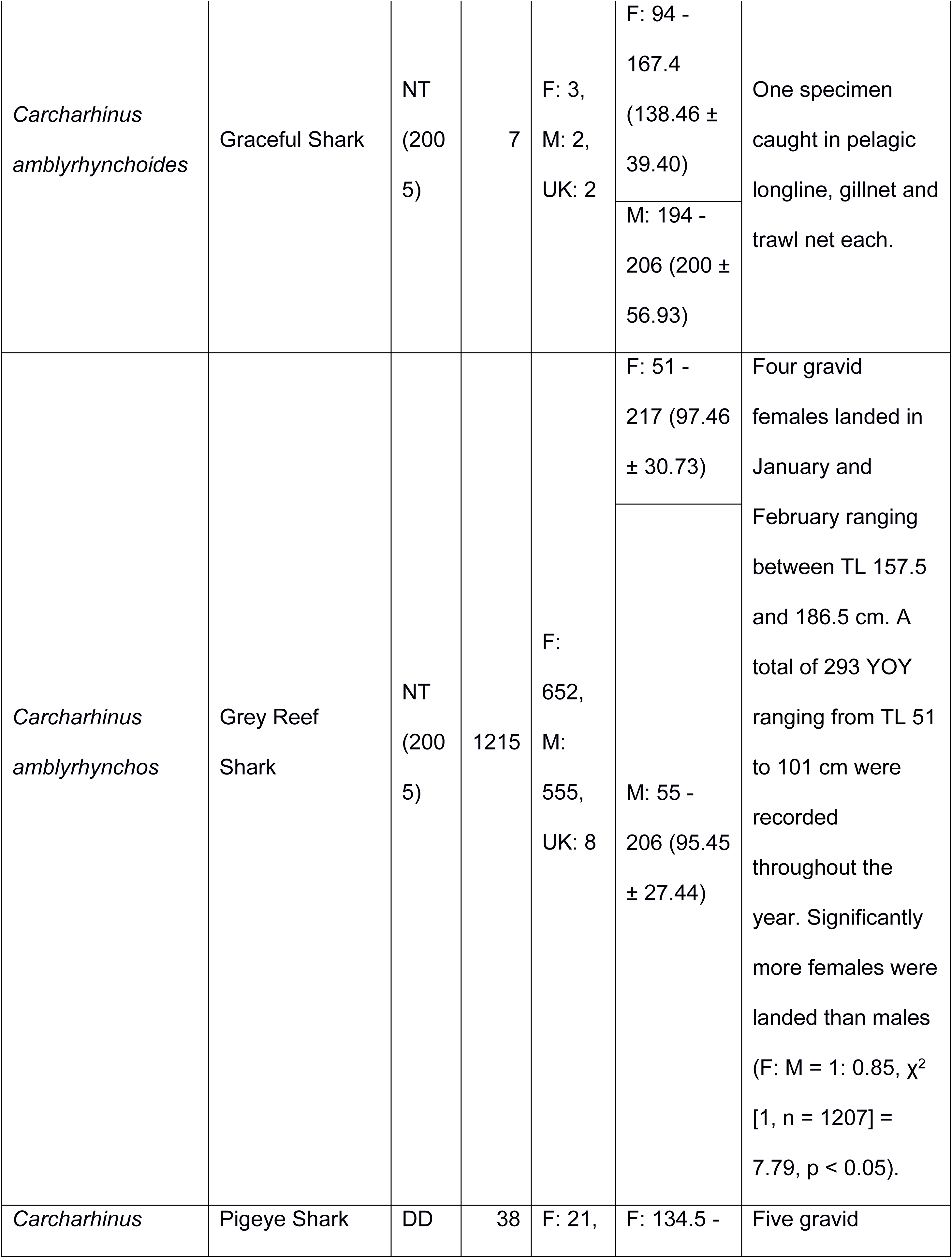
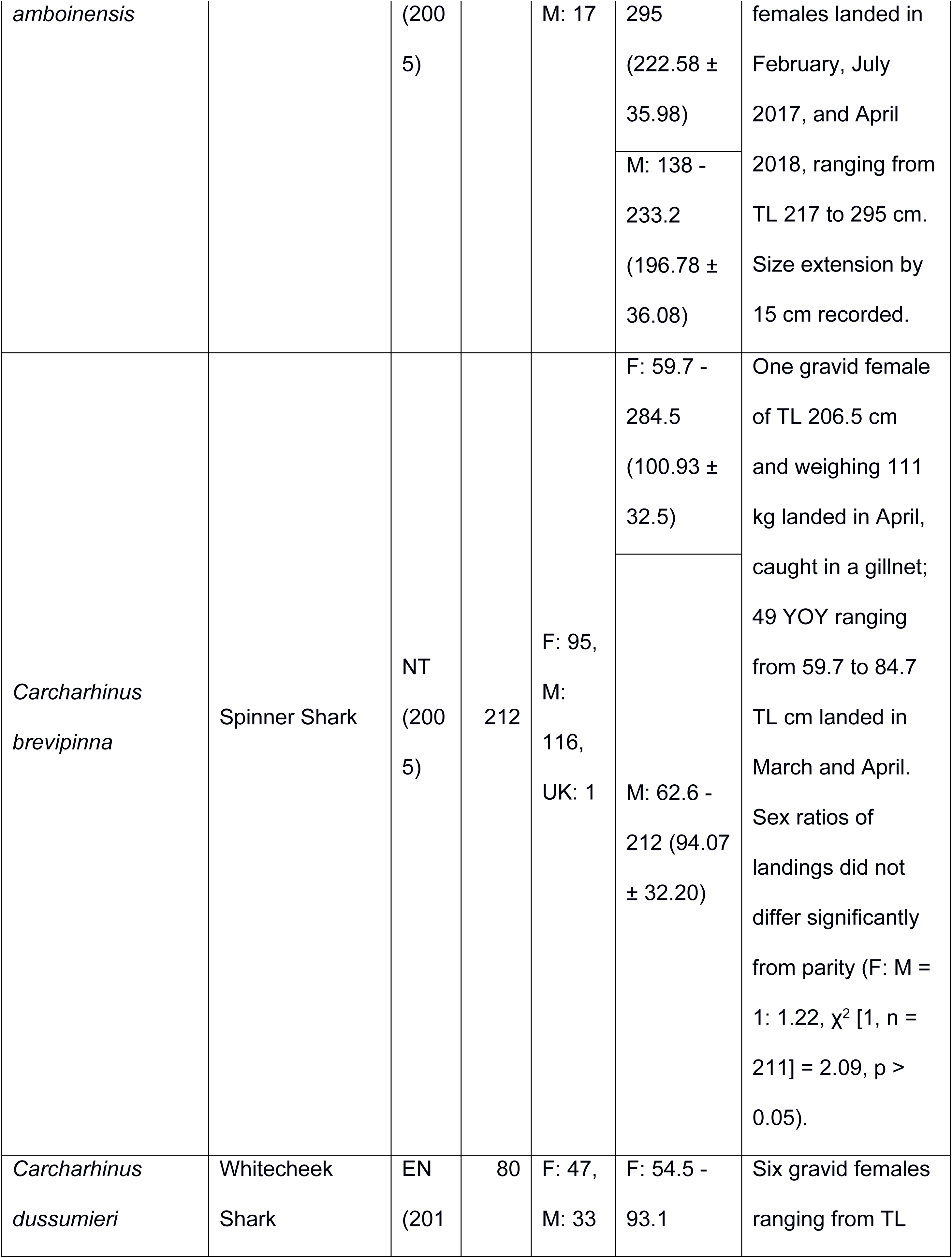
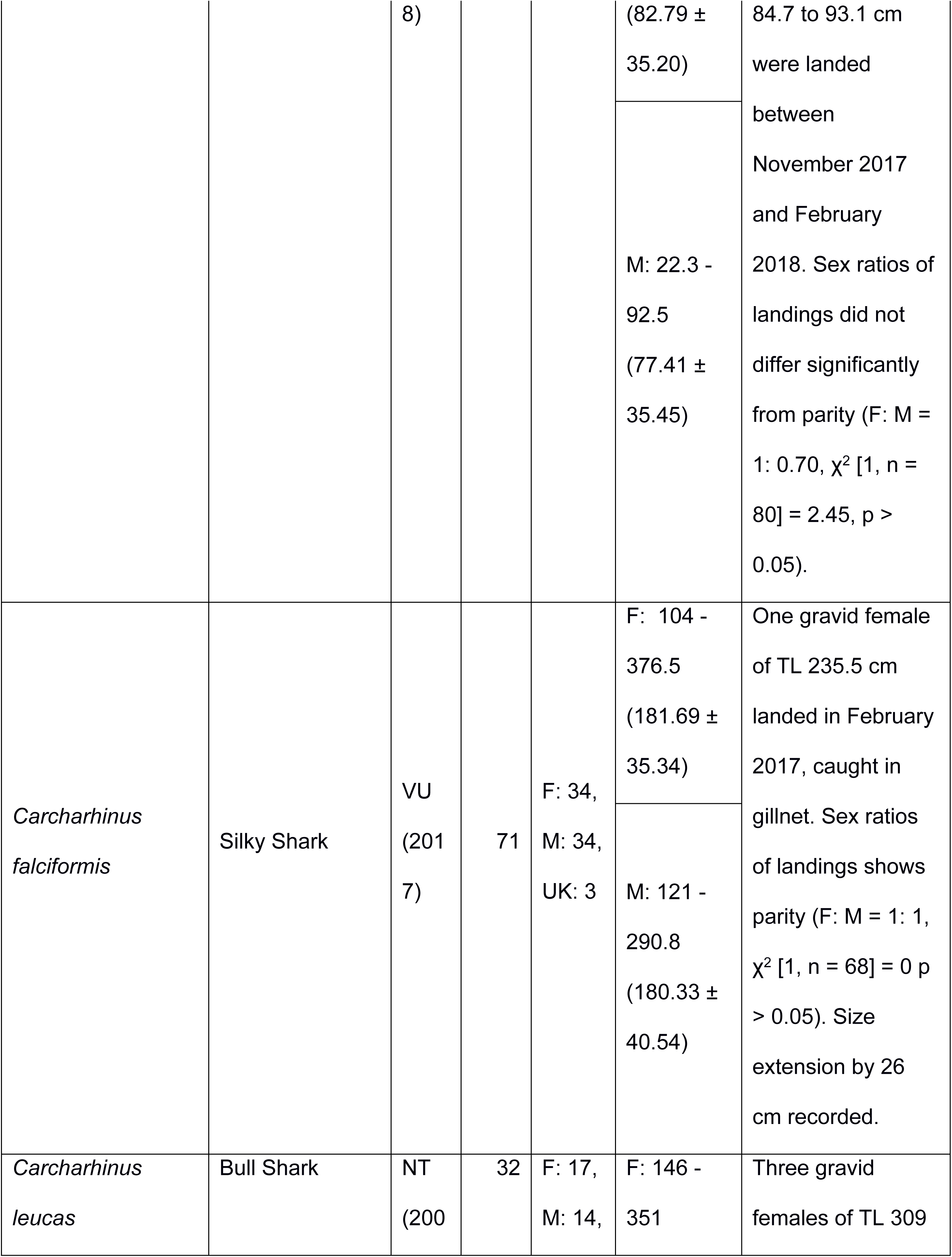
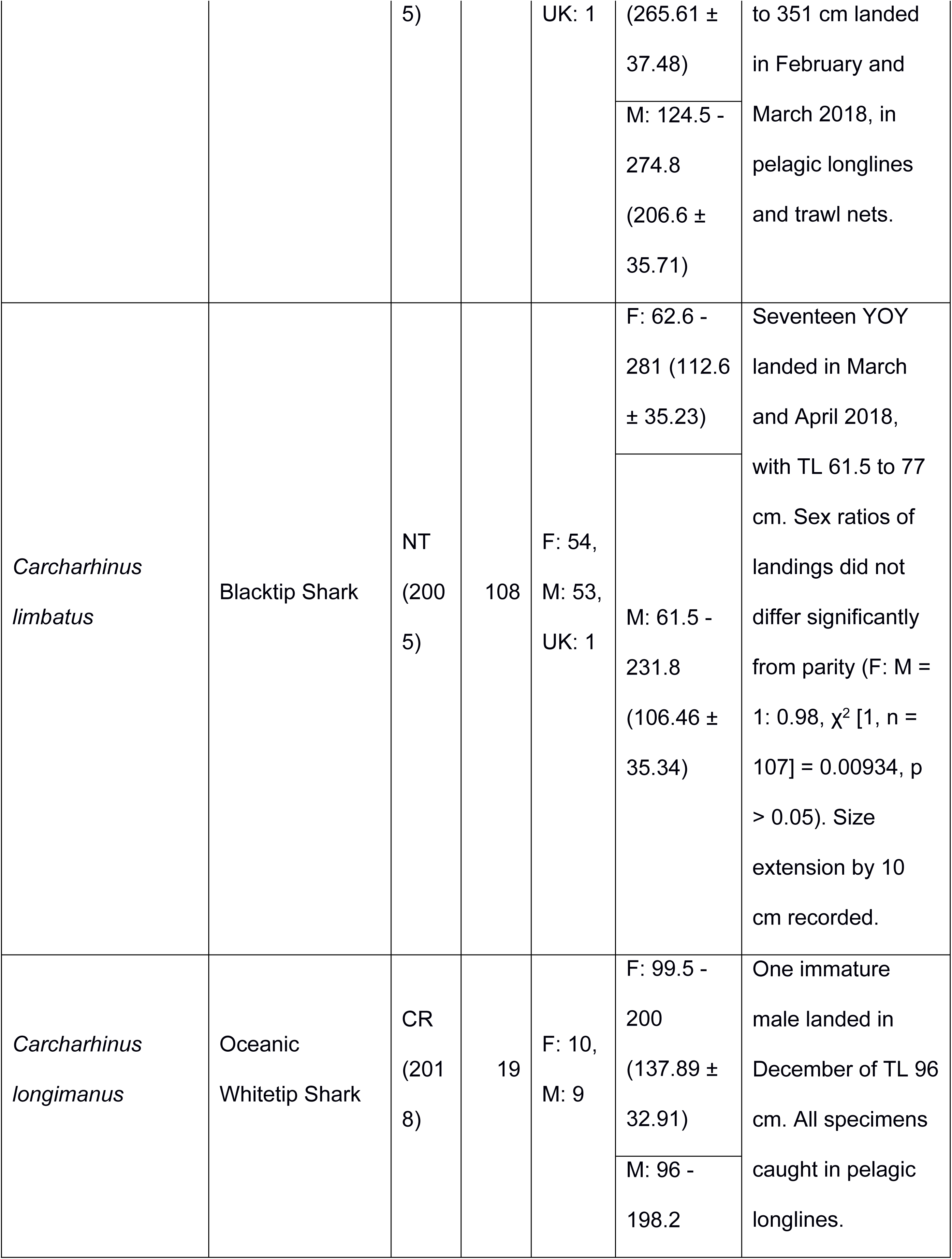
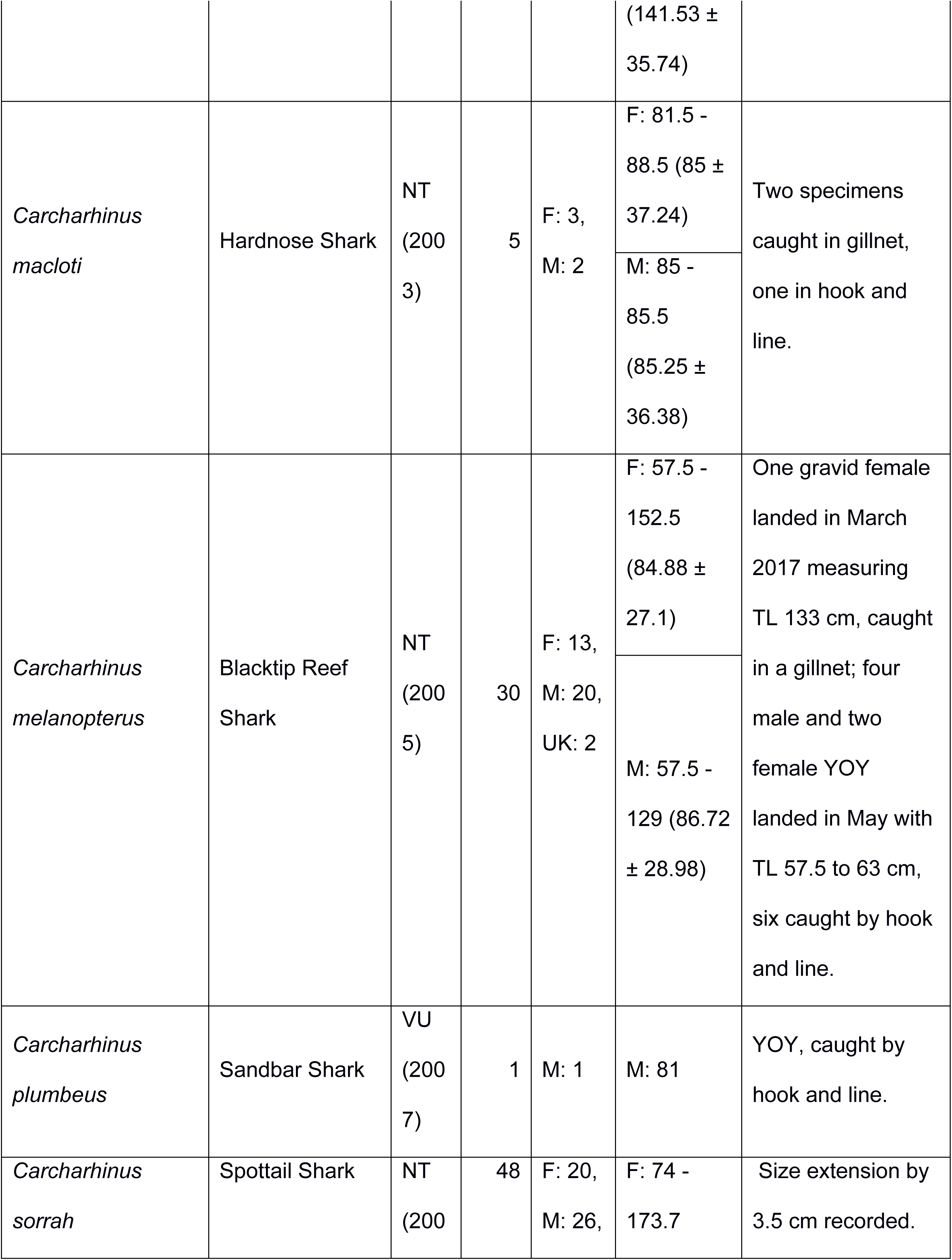
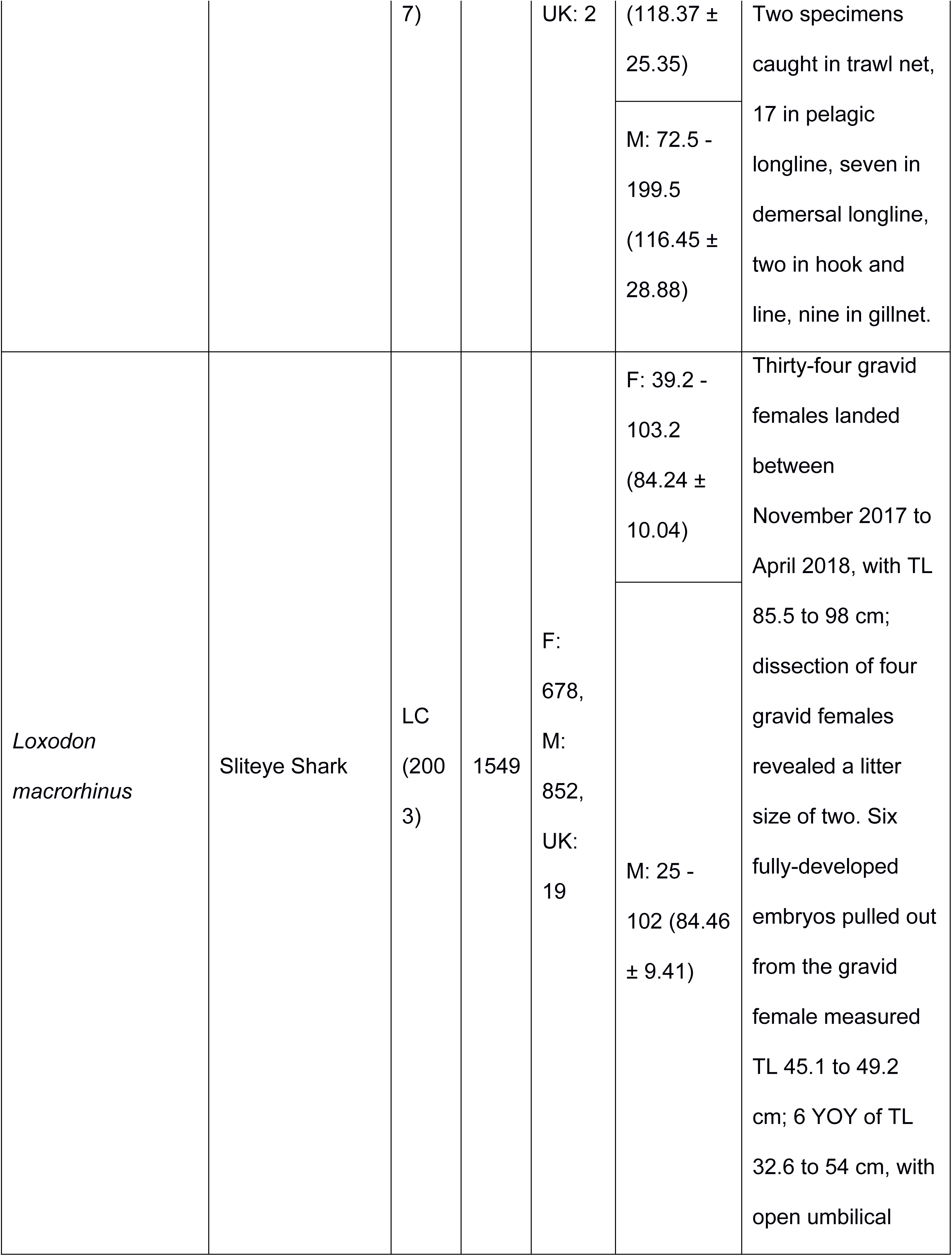
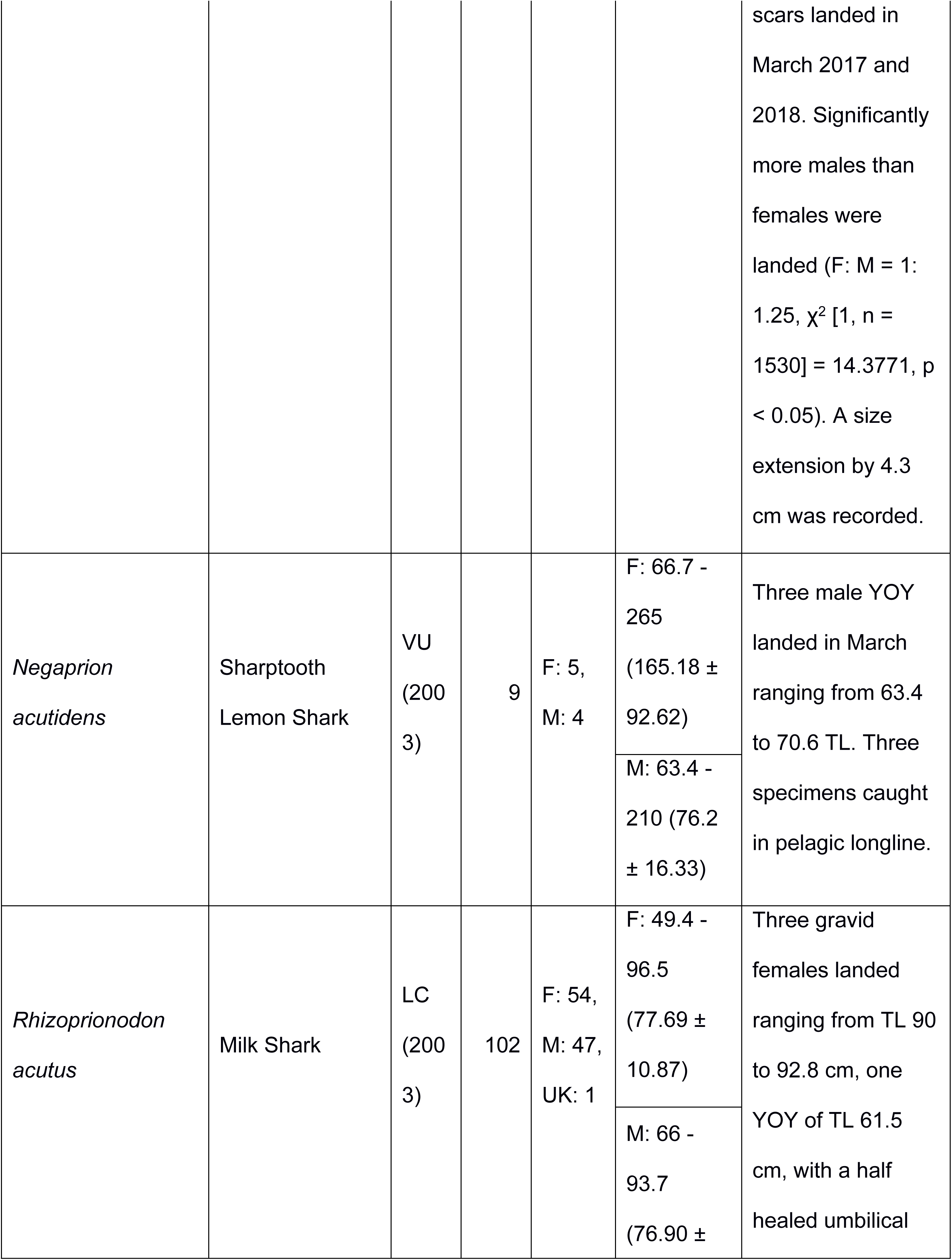
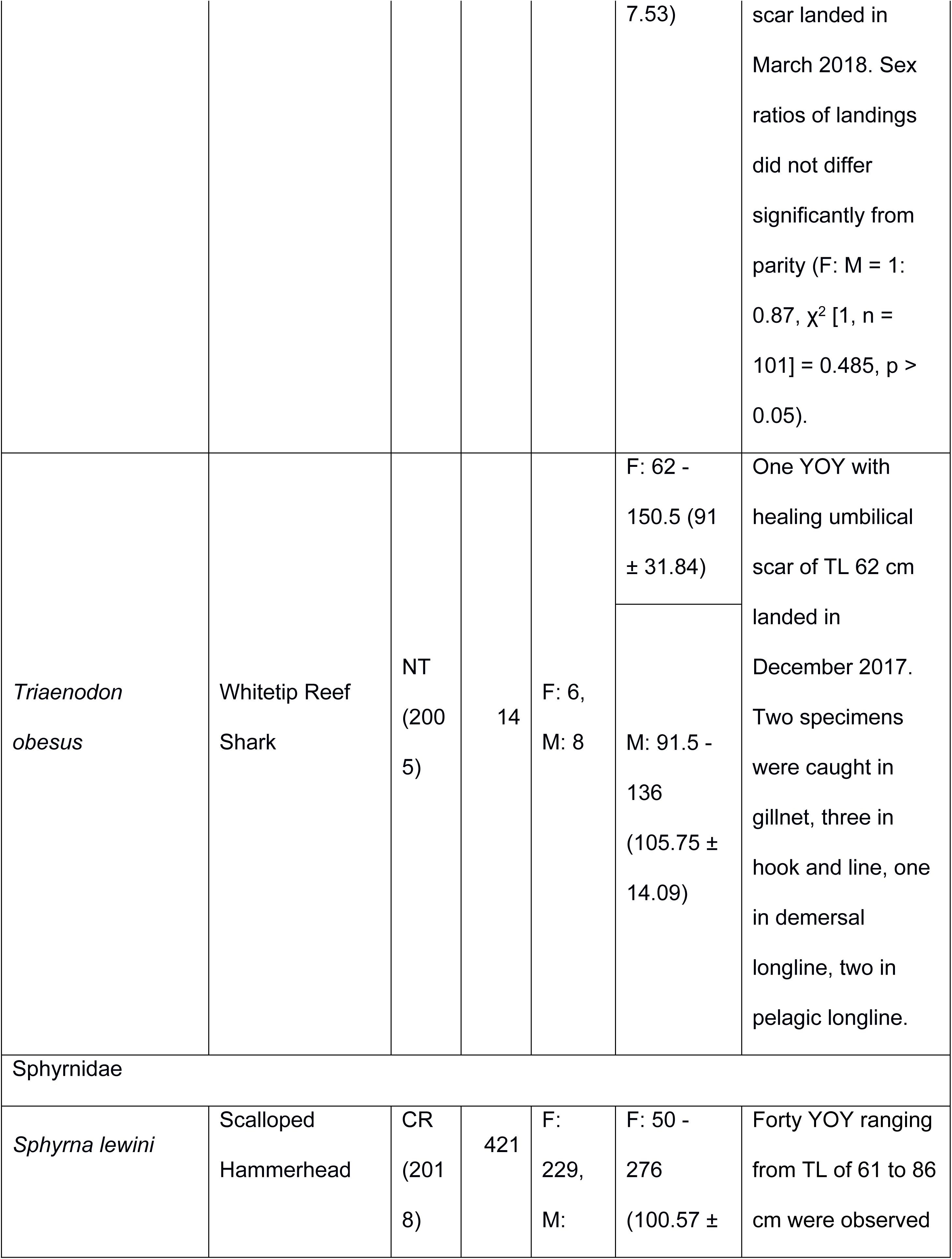
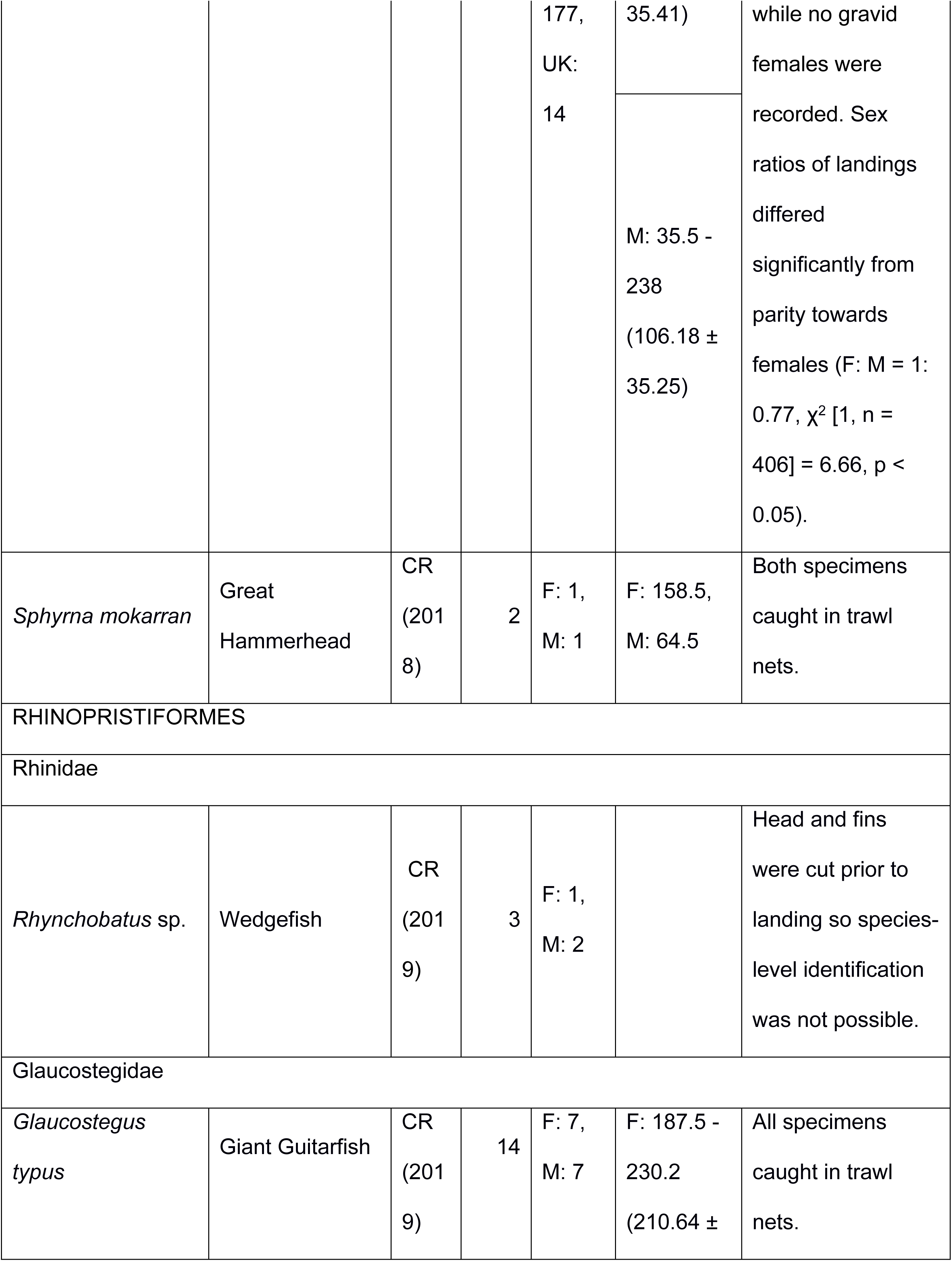
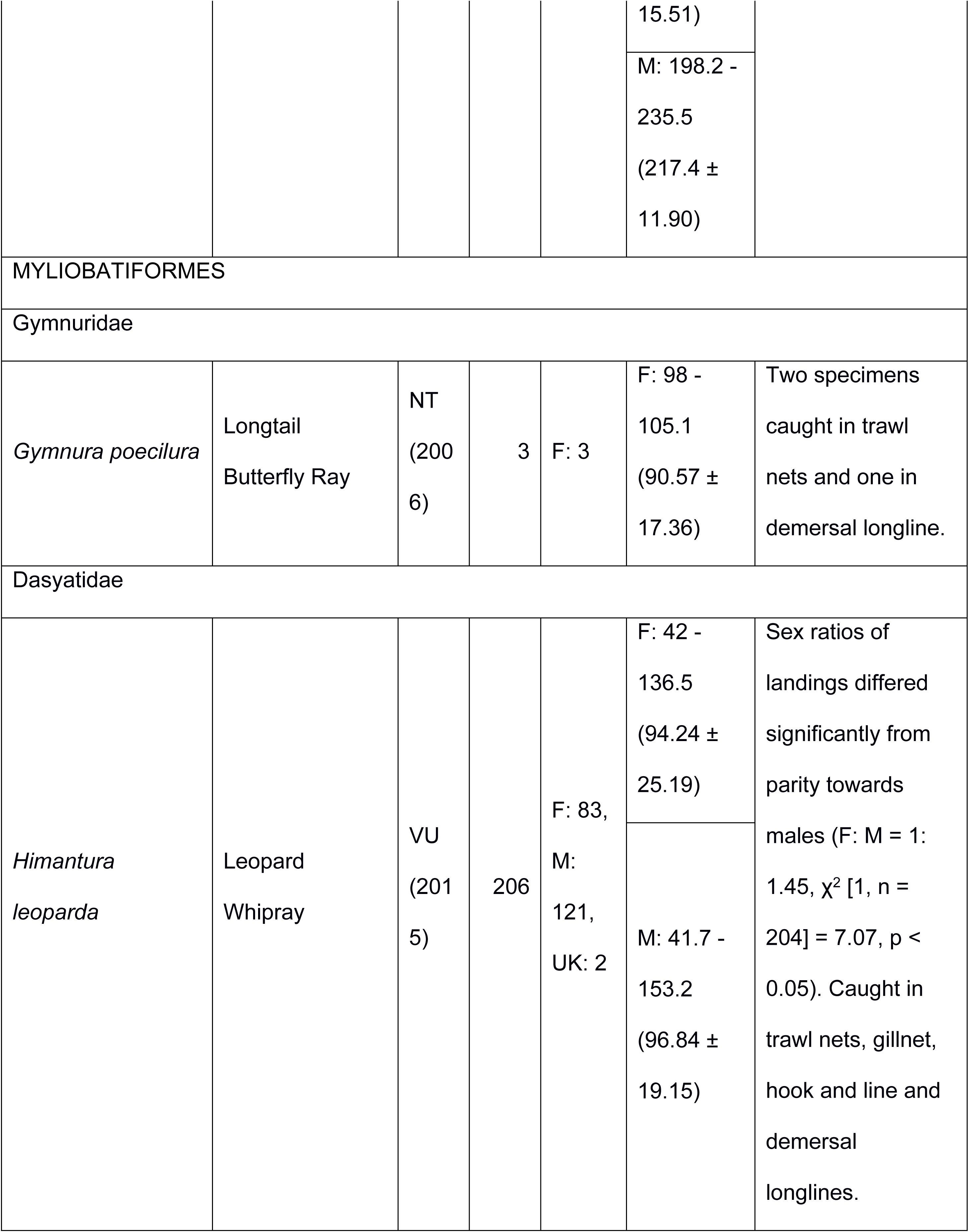
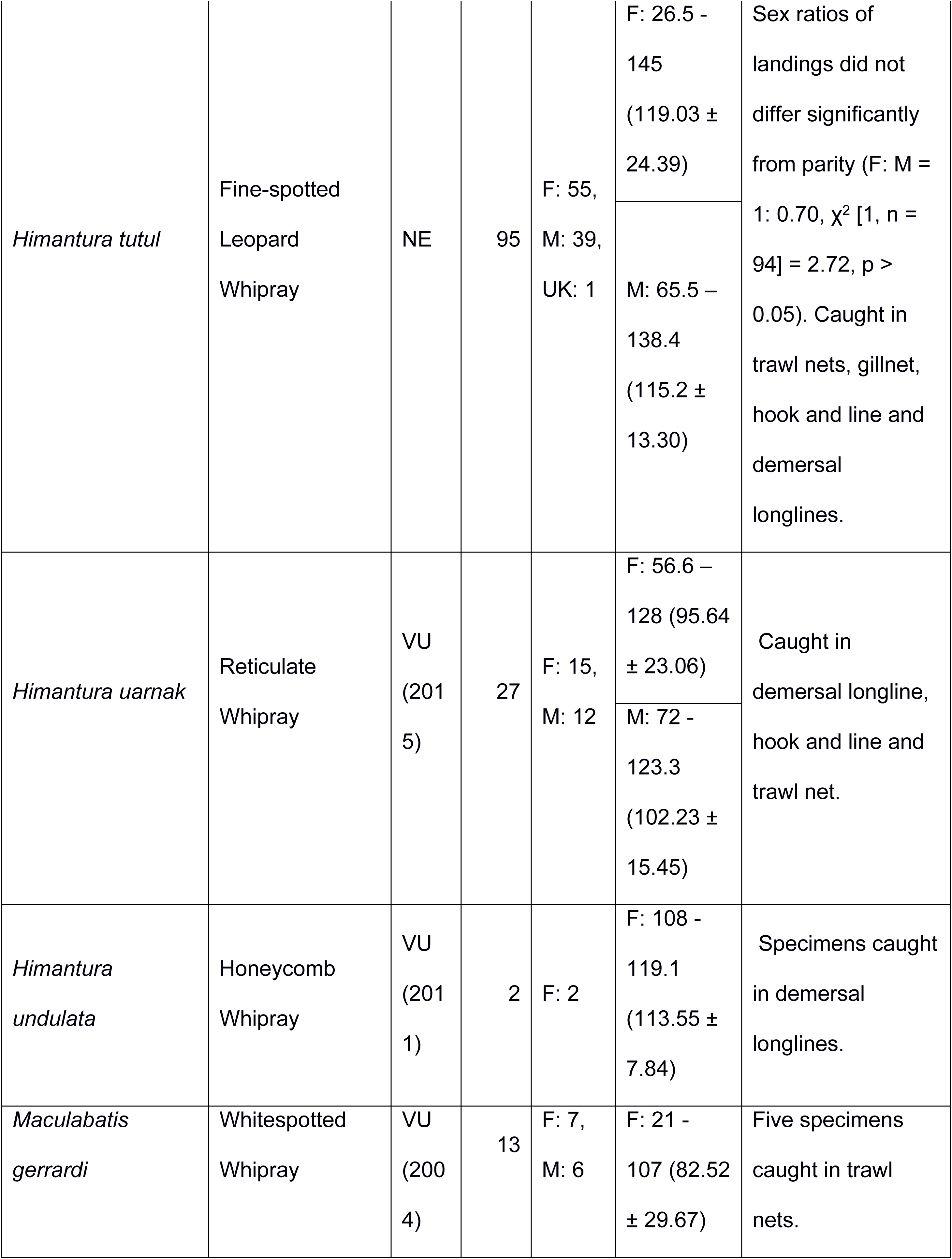
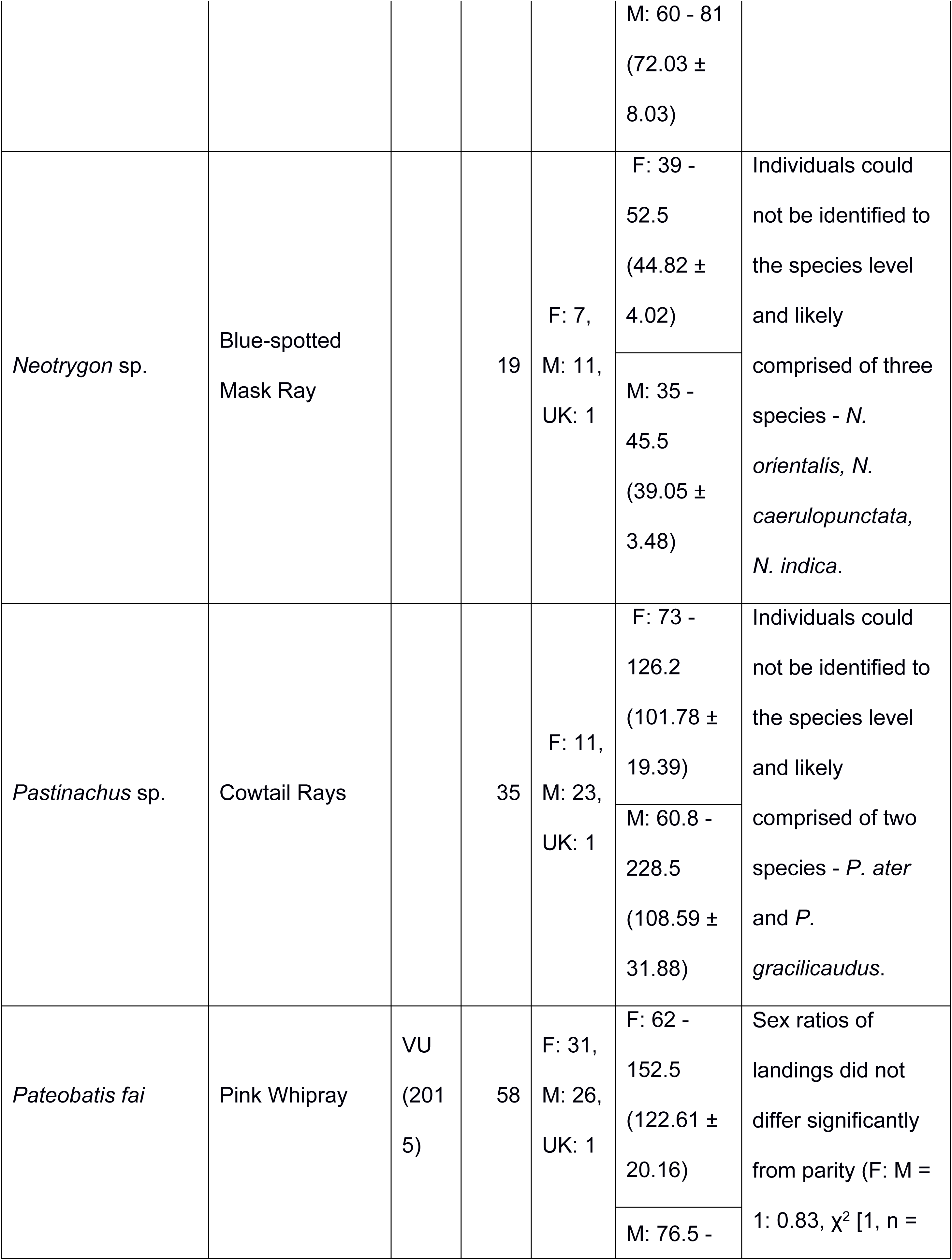
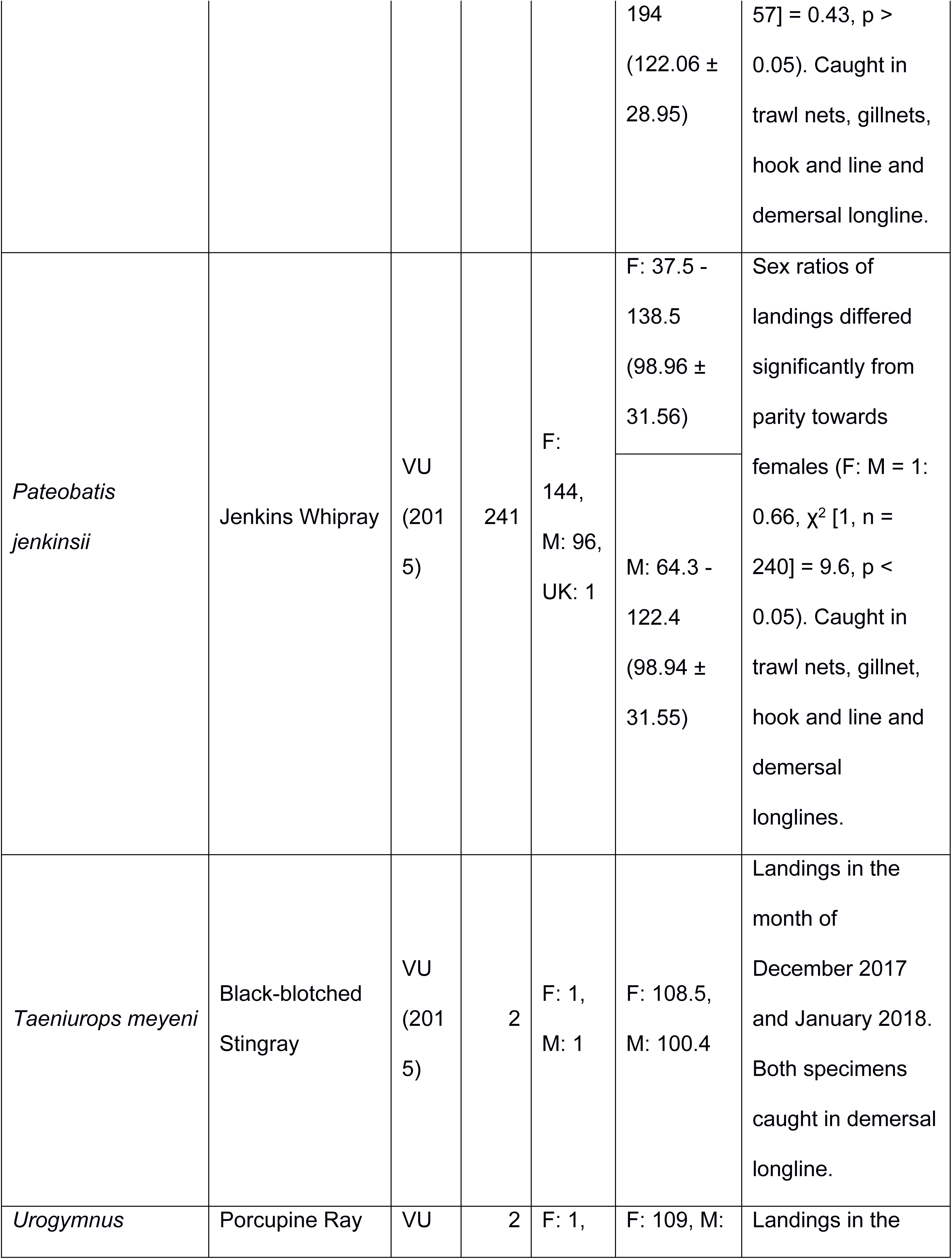
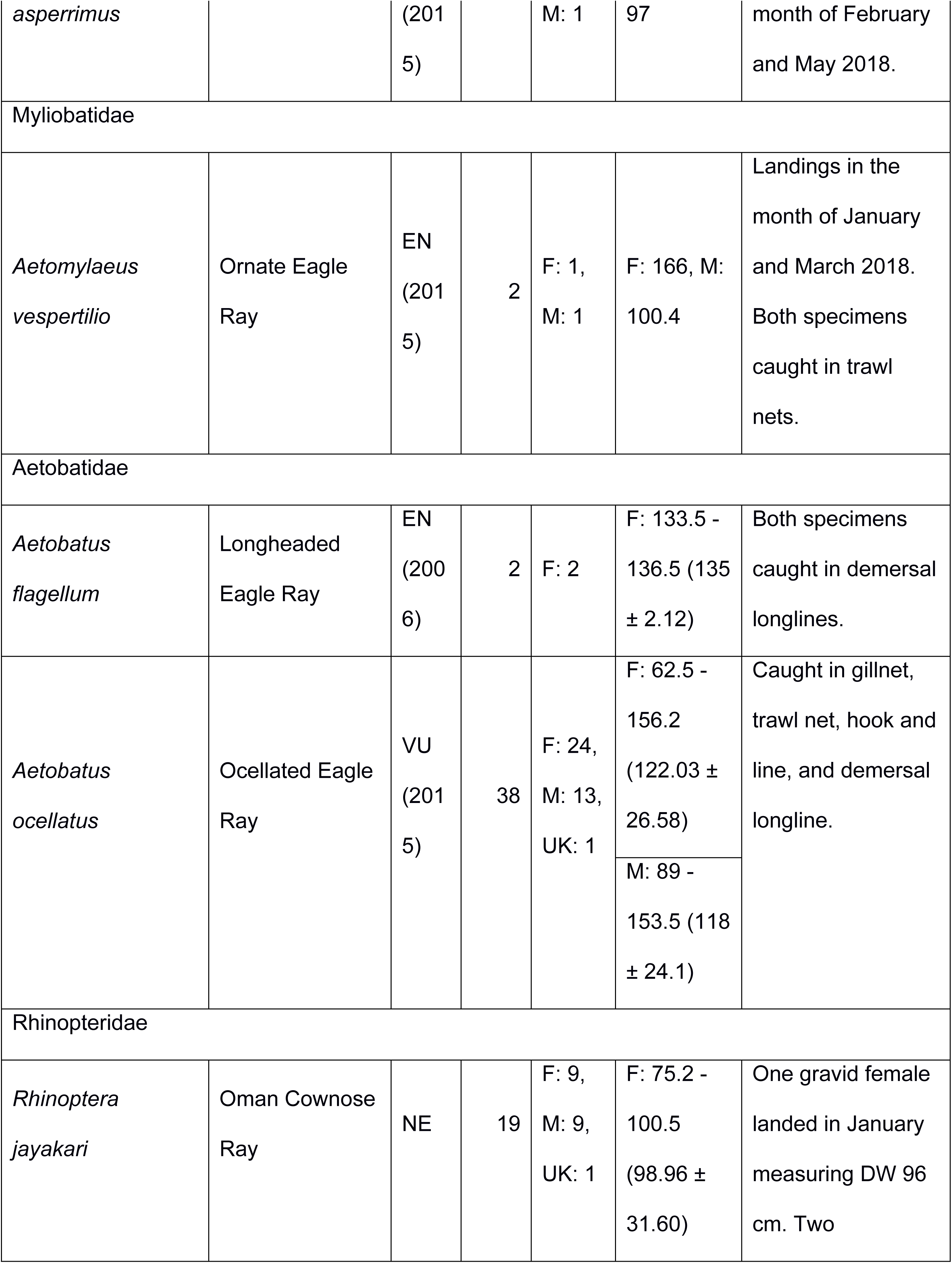
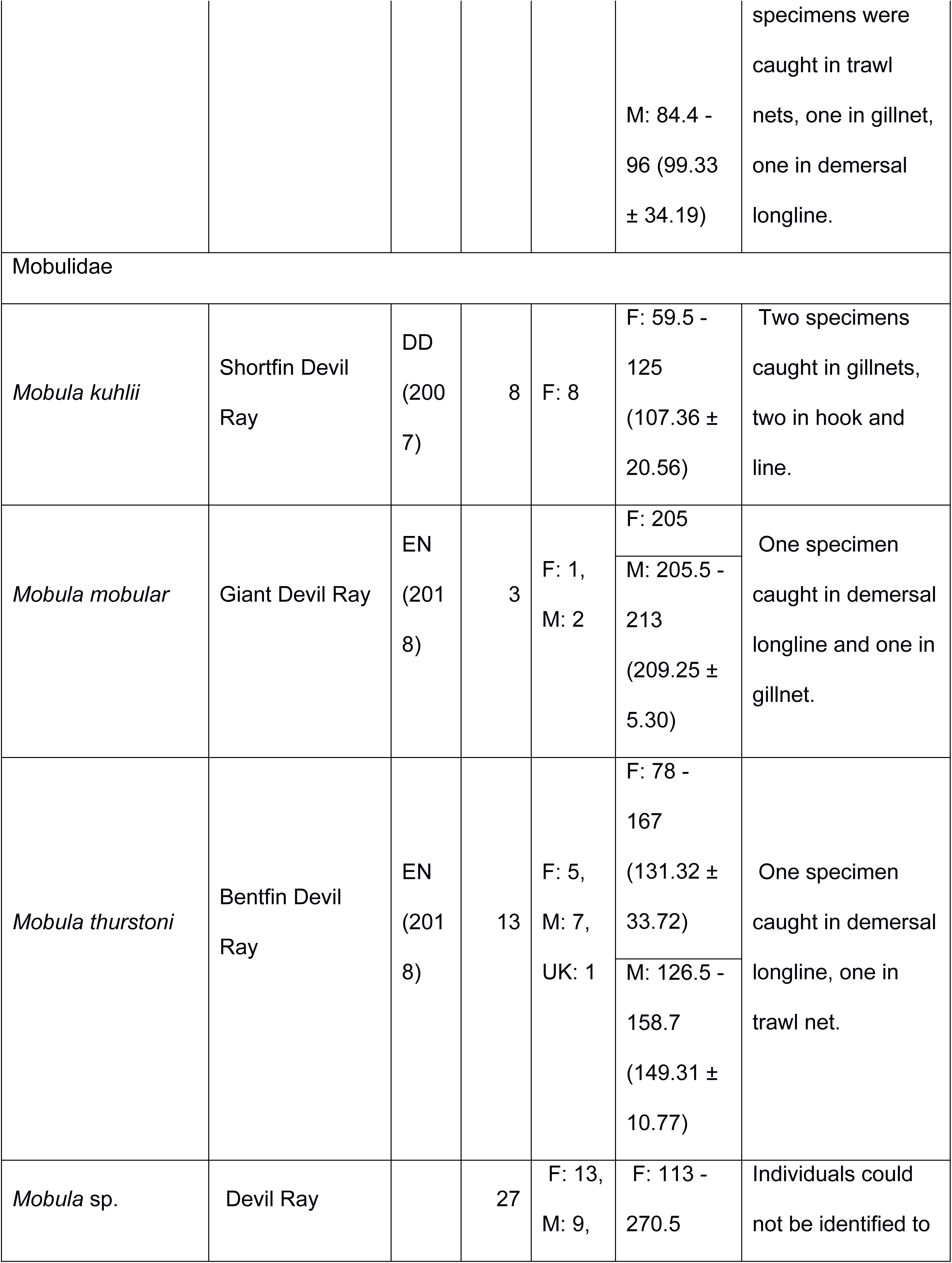
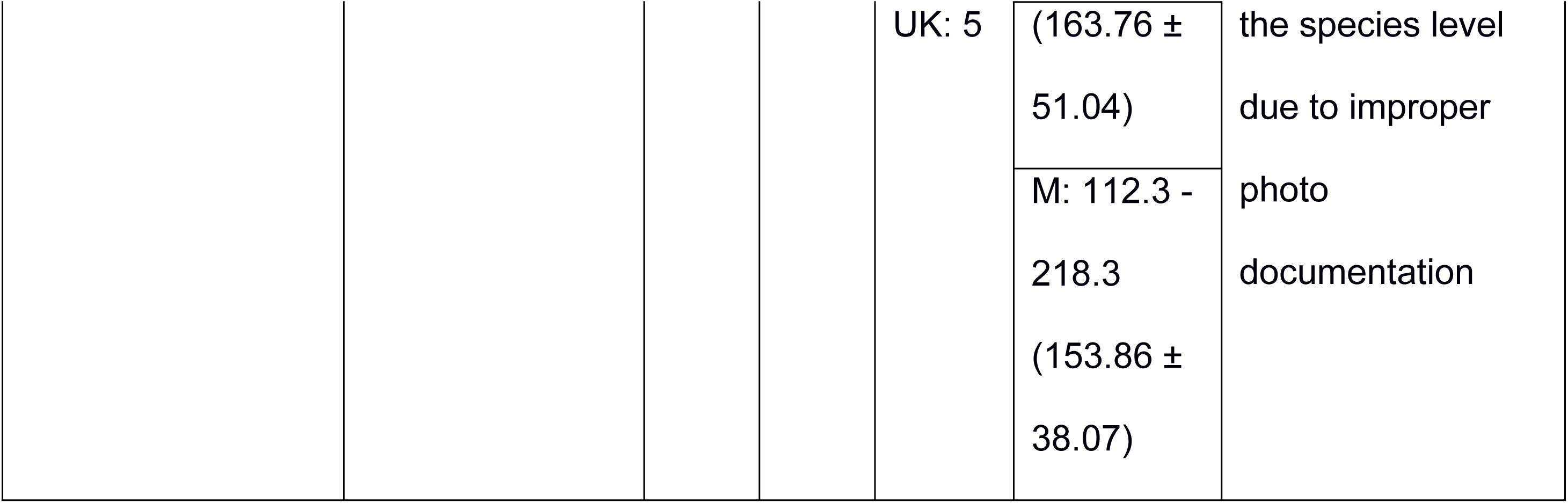
Summary of biological data for sharks and rays landed. The table includes total number of individuals; International Union for Conservation of Nature (IUCN) Red List of Threatened Species status (as of December 2019), where the categories are CR – Critically Endangered, EN – Endangered, VU – Vulnerable, NT – Near Threatened, LC – Least Concern, DD – Data Deficient, NE – Not Evaluated, and year of assessment in parentheses; and sizes (total length for sharks (TL) and disc width (DW) for rays) by sex (F – female; M – male; UK – unknown), presence of gravid females and young of year (YOY). Results of Chi-square tests of parity in sex ratios for shark and ray species are provided for species with ≥ 50 individuals recorded, and fishing gears used to catch the species are provided where applicable.

The next section first provides an overview of the information collected on sharks and then rays separately including species composition, species susceptibility to fishing gear, and biological data of the most abundant species recorded.

## CHONDRICHTHYES: ELAMOBRANCHII: EUSELACHII: SELACHIMORPHA

### Species composition

Species from the Carcharhinidae family dominated landings and accounted for 19 of the 36 species (82.98 %). The six most dominant shark species landed were *Loxodon macrorhinus* (n = 1,549, 33.44 %), *Carcharhinus amblyrhynchos* (n = 1,215, 26.23 %), *Sphyrna lewini* (n = 421, 9.09 %), *C. albimarginatus* (n = 295, 6.36 %), *C. brevipinna* (n = 212, 4.57 %), and *Paragaleus randalii* (n = 169, 3.64 %), constituting 83.35 % of all landings.

The number of sharks sampled across the year ranged from a mean abundance of 41.61 ± 11.58 sharks per day in January to 0.5 ± 0.93 sharks per day in May with landings peaking from November (41.36 ± 11.11 sharks per day) to April (18.90 ± 3.08 sharks per day) (Fig 3).

**Fig 3.**
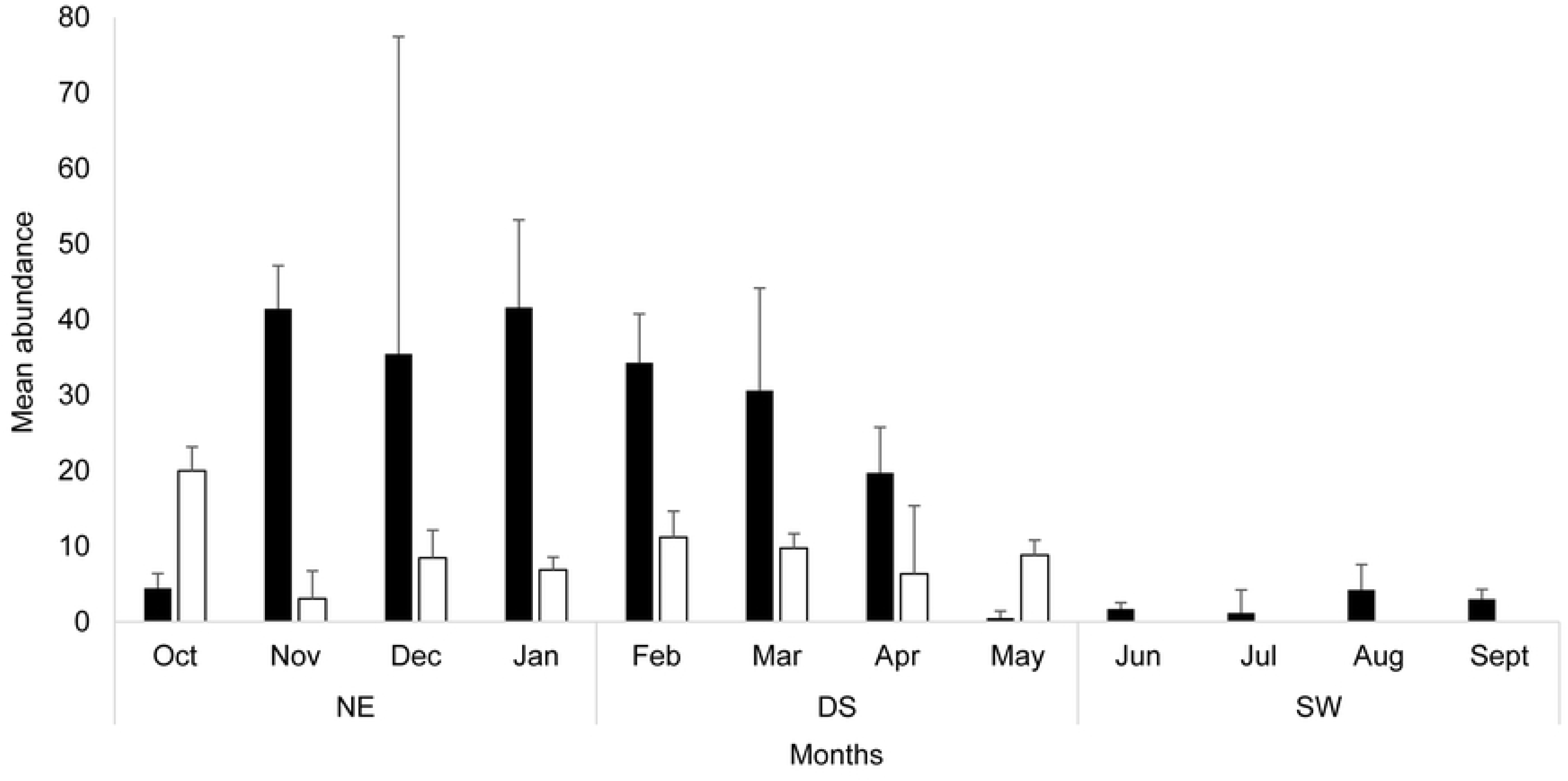
Trends in the mean abundance of daily shark and ray individuals landed across months. Shark abundance is represented in black, and rays in white. The seasons are north-east monsoon (NE) (October–January), dry season (DS) (February–May) and south-west monsoon (SW) (June–September). Error bars indicate standard error.

### Use of fishing gears and fishing grounds

Twenty-one species were recorded interacting with gillnets, hook and line, and pelagic longlines each, 18 species were recorded interacting with demersal longlines, 14 species with trawl nets and two species (*Centrophorus atromarginatus* (n=1) and *C. granulosus* (n = 6)) with deep-sea longlines.

Certain species were only recorded in one type of gear. For example, *Alopias pelagicus* (n = 28), *A. superciliosus* (n = 6), *C. longimanus* (n = 19) and *Hemitriakis indroyonoi* (n = 2) were only associated with pelagic longlines; *Mustelus mosis* (n = 7) were only recorded from hook and line; and *S. mokarran* (n = 2) were only recorded in trawl nets.

Further, there was a significant difference between the TL of sharks caught depending on the type of fishing gears used (*f* (5, 2,146) = 88.66, *p* < 0.005). Sharks landed in pelagic longliners had a high TL range from 21.5 to 376.5 cm (mean of 124.90 ± 49.83); those in demersal longlines had a TL range from 42 to 214.5 cm (mean 18.81 ± 93.76); those in deep-sea longlines (>200 m) had a TL range from 72.5 to 103 cm (mean of 88.3 ± 10.80); those in gillnets had a TL range from 25 to 312.5 cm (mean of 97.49 ± 34.26); those in trawl nets had a TL range from 50 to 297.9 cm (mean of 47.67 ± 97.65); and those from hook and line had a TL range of 46 to 266.7 cm (mean of 47.67 ± 97.65) (Fig 4). The fishing grounds with frequency of each fishing gear used across the islands is provided in Fig 1.

**Fig 4.**
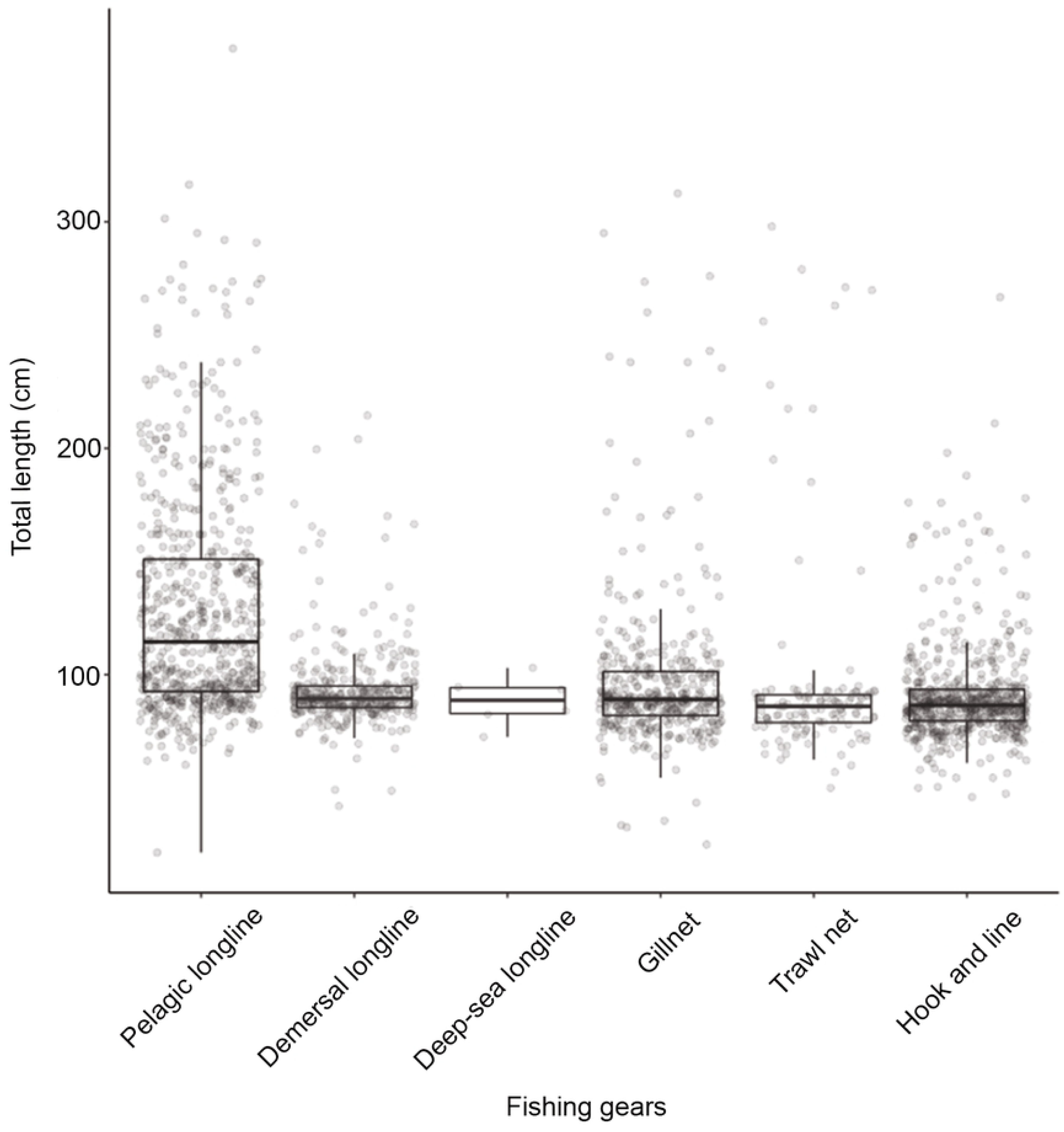
Total length (in cm) of sharks landed across the different fishing gear used on the islands.

### Seasonality, size frequency, and length-weight relationships

The following section provides details of the size frequency, seasonality and length-weight relationships of the six dominant shark species landed. Additional information on all species, including sex ratios where applicable and recorded size extensions for seven species, are provided in Table 1. For the non-dominant shark species in landings, of the 2,258 male individuals whose maturity was recorded, 35.93 % of sharks were mature. The majority of specimens from small-bodied species (TL < 1.5 m) were mature (63.48%) whereas the majority of specimens from large-bodied species (TL > 1.5 m) were immature (84.79%).

### CARCHARHINIFORMES - CARCHARHINIDAE - *Loxodon macrorhinus*

The size frequency of *L. macrorhinus* followed a unimodal size distribution where mature individuals of TL 85 - 95 cm (n = 830, 54.35 %) were dominantly landed across both sexes (Fig 5). Landings were variable across seasons with a peak during the dry season (n = 909) followed by NE monsoon (n = 632) and low landings during the SW monsoon (n = 8) (Fig 6).

**Fig 5.**
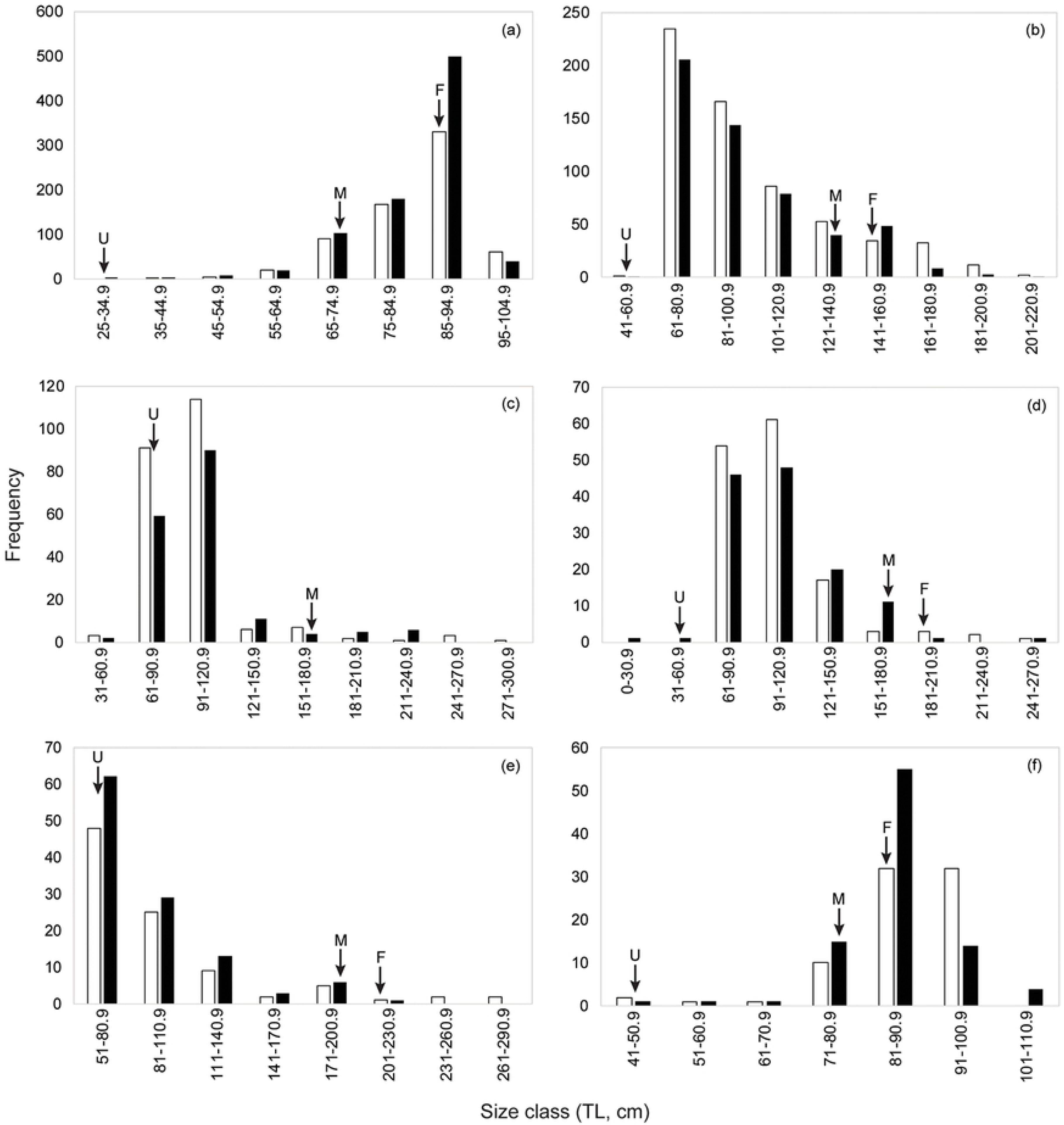
Size frequency distribution for males and females for the six most commonly landed shark species. (a) *Loxodon macrorhinus*, (b) *Carcharhinus amblyrhynchos*, (c) *Sphyrna lewini,* (d) *Carcharhinus albimarginatus,* (e) *Carcharhinus brevipinna,* and (f) *Paragaleus randalli*. The black bars represent males and the white bars represent females. The arrows represent the smallest individual representing young of year with the presence of an umbilical scar ‘U’, ‘F’ the smallest gravid females recorded, and ‘M’ the smallest recorded mature males.

**Fig 6.**
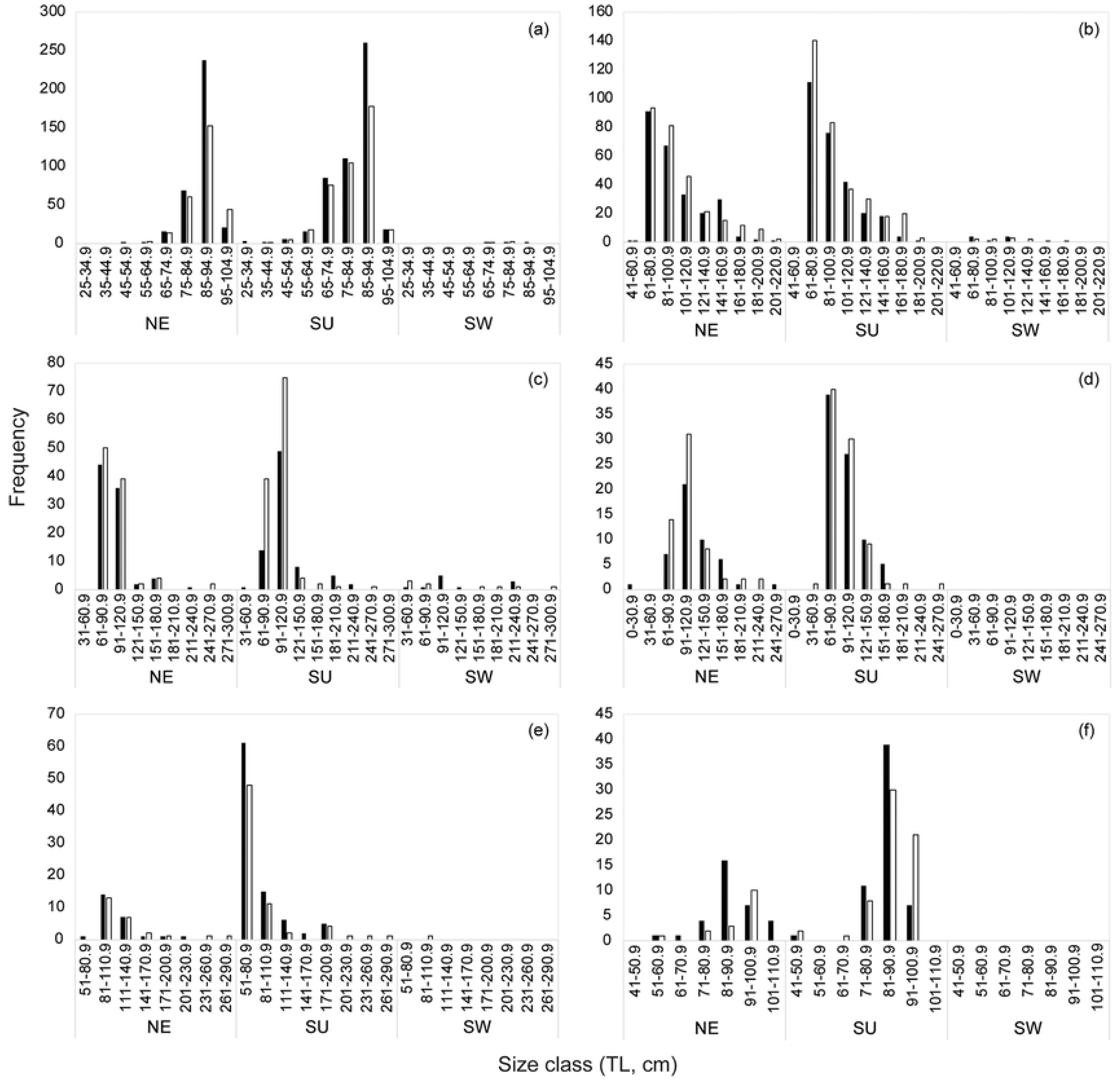
The seasonal size distribution of male and females for the six most commonly landed shark species. (a) *Loxodon macrorhinus*, (b) *Carcharhinus amblyrhynchos*, (c) *Sphyrna lewini,* (d) *Carcharhinus albimarginatus,* (e) *Carcharhinus brevipinna,* and (f) *Paragaleus randalli.* The seasons are north-east monsoon (NE) (October–January), dry season (DS) (February–May) and south-west monsoon (SW) (June–September). The black bars represent males and the white bars represent females.

Of the 852 males, 75.94 % were mature. The smallest immature male was 32.6 cm whereas the largest was 78.1 cm. The smallest mature male was 67.3 cm, whereas the largest was 102 cm with a TL_50_ of 70.61 cm (Fig 7). Landings of gravid females at various stages of embryo development were observed throughout the year, whereas YOY were observed in the month of March and April 2017 and 2018, with one individual observed in January 2018.

**Fig 7.**
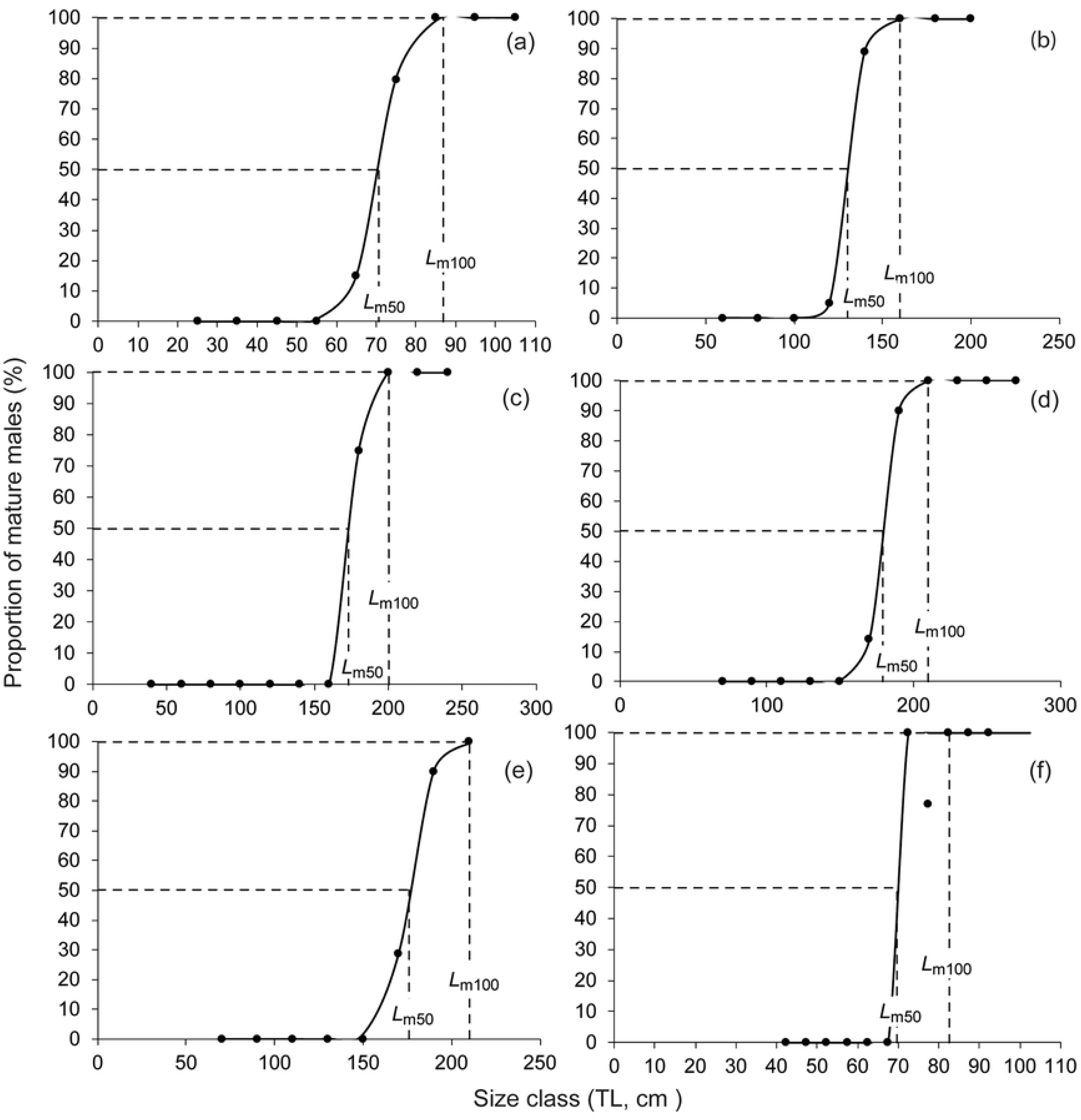
Percentage of mature males with total length (TL) for sharks at 50% and 100% maturity for the six most commonly landed shark species. (a) *Loxodon macrorhinus* (n = 820), (b) *Carcharhinus amblyrhynchos* (n = 518), (c) *Sphyrna lewini* (n = 176), (d) *Carcharhinus albimarginatus* (n = 124), (e) *Carcharhinus brevipinna* (n = 87), and (f) *Paragaleus randalli* (n = 91)

The length-weight relationships differed between sexes, where females showed positive allometry (b = 3.40), whereas for males, the weight increased in an almost allometric manner (b = 2.99), in proportion with the cube of the length (Fig 8, S1 Table).

**Fig 8.**
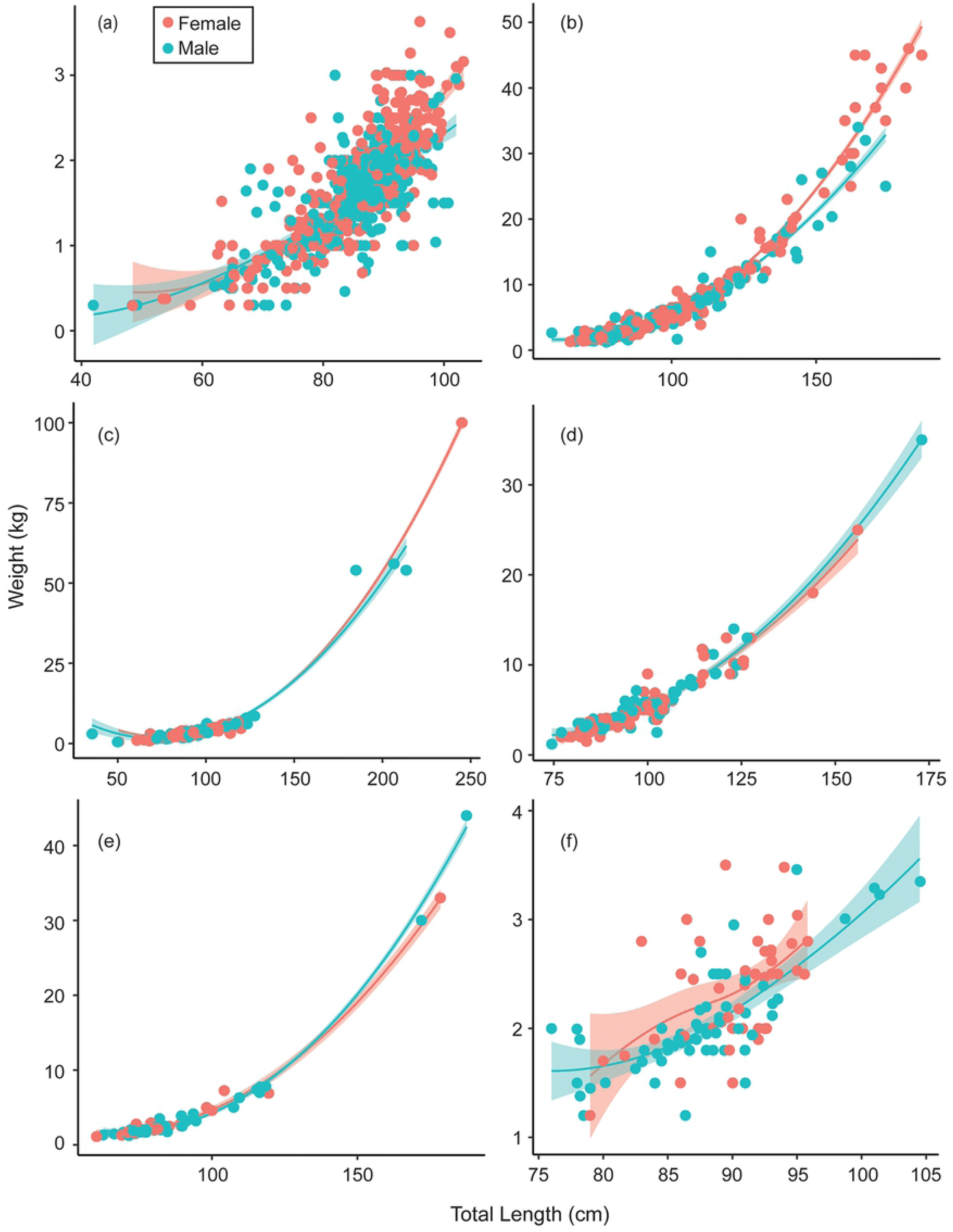
Length and weight relationships between total body mass (kg) and total length (cm) for the six most commonly landed shark species. (a) *Loxodon macrorhinus*, (b) *Carcharhinus amblyrhynchos*, (c) *Sphyrna lewini,* (d) *Carcharhinus albimarginatus,* (e) *Carcharhinus brevipinna,* and (f) *Paragaleus randalli*.

### CARCHARHINIFORMES - CARCHARHINIDAE - *Carcharhinus amblyrhynchos*

Immature individuals of size class TL 61 - 81 cm dominated landings across both sexes (n = 441, 38.28 %), followed by size class 81 - 100.9 cm (n = 310, 26.90 %) (Fig 5). Landings were variable across seasons with a peak during the dry season (n = 633) followed by NE monsoon (n = 559) and a lower number of individuals landed during the SW monsoon (n = 23) (Fig 6).

Of the 555 males, 16.19 % were mature. The smallest mature male was TL 126.3 cm whereas the largest was 206 cm. The TL_50_ of males was 131.69 cm (Fig 7).

The length-weight relationships did not differ between sexes, where both the sexes showed a positive allometric relationship (female b = 3.45; male b = 3.29), in proportion with the cube of the length (Fig 8, S1 Table).

### CARCHARHINIFORMES - SPHYRNIDAE- *Sphyrna lewini*

Landings of *S. lewini* were dominated by the size class TL 91 to 120.9 cm (n = 204, 50.37 %), followed by size class 61 - 90.9 cm (n = 150, 37.03 %) (Fig 5). Landings were variable across seasons with a peak during the dry season (n = 211) followed by NE monsoon (n = 189) whereas comparatively fewer landings were recorded during the SW monsoon (n = 21) (Fig 6).

Of 177 males, 9.65 % were mature. Immature individuals measured 35.5 to 170.4 cm TL, whereas mature individuals measured 177 to 238 cm TL with a TL_50_ of 173 cm (Fig 7).

The length-weight relationships did not differ between sexes, where both the sexes showed a positive allometric relationship (female b = 2.91; male b = 2.60), in proportion with the cube of the length (Fig 8, S1 Table).

### CARCHARHINIFORMES - CARCHARHINIDAE - *Carcharhinus albimarginatus*

The size frequency of *C. albimarginatus* followed a unimodal size distribution where immature individuals of size class TL 91 - 121 cm dominated landings (n = 109, 40.37 %) across both sexes (Fig 5). Landings were variable across seasons with a peak during the dry season (n = 177), followed by NE monsoon (n = 118) with none recorded during the SW monsoons (Fig 6).

Of the 137 males, 4.47 % were mature. The smallest mature male was 173 cm whereas the largest was 249 cm. The TL_50_ of males was 178.98 cm (Fig 7).

The length-weight relationships showed that males and females did not differ significantly in their average weight for a given length, and weight increased in a positive allometric manner (female b = 3.65; male b = 3.55), in proportion with the cube of the length (Fig 8, S1 Table).

### CARCHARHINIFORMES - CARCHARHINIDAE - *Carcharhinus brevipinna*

Juveniles of the size class TL 51 - 80.9 cm (n = 110, 52.88 %) dominated landings, followed by size class 81 - 110.9 cm (n = 54, 25.96 %) where male YOY (n = 62) were more abundant than females (n = 48) (Fig 5). Landings were variable across seasons and differed in sex and size. Landings peaked during the dry season (n = 159) followed by NE monsoon (n = 52) with low landings during the SW monsoon (n = 1) (Fig 6).

Of the 116 males sampled, 10.11 % were mature. Mature males ranged from TL 172 to 212 cm, whereas immature males ranged from TL 62.6 to 175.78 cm. The TL_50_ of males was 175.78 cm (Fig 7).

The length-weight relationships differed between sexes, where the female showed positive allometry (b = 3.20), whereas for males, the weight increased in a near perfectly allometric manner (b = 3.23), in proportion with the cube of the length (Fig 8, S1 Table).

### CARCHARHINIFORMES - HEMIGALEIDAE - *Paragaleus randalli*

The size frequency followed a unimodal distribution where females of size classes TL 81 - 90.9 cm (n = 87, 51.47 %) dominated landings, followed by 91 - 100.9 cm (n = 46, 27.2 %) (Fig 5). Landings peaked during the dry season (n = 120) followed by a decrease in NE monsoon (n = 49) whereas no landings were observed during the SW monsoons (Fig 6).

Of 91 males recoded, 93.4 % were mature. The smallest immature individual measured 43.5 cm whereas the largest measured 76.5 cm. The smallest mature individual measured 74.5 cm, whereas the largest measured 106.2 cm with a TL_50_ of 69.6 cm (Fig 7).

The length-weight relationships showed that males and females did not differ significantly in their average weight for a given length, and weight increased in a positive allometric manner (female b = 2.65; male b = 2.59), in proportion with the cube of the length (Fig 8, S1 Table).

## RAYS

### CHONDRICHTHYES: ELASMOBRANCHII: BATOIDEA

#### Species composition

Species from the *Dasyatidae* family dominated landings, accounting for 11 of the 21 species, and 63.06 % of the total landings. The three most common rays landed were *Pateobatis jenkinsii* (n = 241, 21.71 %), *Himantura leopard*a (n = 206, 18.55 %) and *H. tutul* (n = 95, 8.55 %); representing 48.82 % of the total ray landings.

The number of rays sampled across the year ranged from 11.2 ± 3.45 rays per day in February to 3.07 ± 3.66 rays per day in November with no pattern observed in landings (Fig 3).

#### New species records

Three species of rays, *Aetobatus flagellum*, *H. tutul*, and *P. fai*, were recorded for the first time from the Andaman and Nicobar Islands (Table 1, S1 Fig).

Eight species previously not confirmed but reported as possibly occurring on the islands by Kumar et al. [24] have been confirmed: *Aetomylaeus vespertilio, Glaucostegus typus, H. undulata, Mobula kuhlii*, *M. tarapacana, Pastinachus ater*, *P. jenkinsii*, and *Urogymnus asperrimus*.

#### Use of fishing gears and fishing grounds

Sixteen species of rays were captured in demersal longlines, 14 species in trawl nets, ten in gill nets, seven in hook and line, and two in pelagic longlines. No rays were captured in deep-sea longlines. Certain species were caught exclusively in certain gears. For example, *A. vespertilio* (n = 2), and *Maculabatis gerrardi* (n = 13) were only caught in trawl nets.

#### Biological traits for rays

Of the 513 male individuals recorded, 80.71 % were mature. Sex ratios were calculated for four rays, *H. tutul*, *H. leoparda*, *P. fai* and *P. jenkinsii* (Table 1).

## DISCUSSION

This is the first systematic landing survey carried out for sharks and rays in the Andaman and Nicobar Islands, contributing to the current knowledge of species diversity and biology for the south and south-east Asian region. Three ray species are new records for the Andaman and Nicobar Islands, including one new record for India, increasing the elasmobranch diversity for the Andaman and Nicobar Islands from 103 to 106, and for India to 152 [24]. A threshold was reached in terms of shark species recorded, but additional efforts are required to fully document ray diversity. The diversity and high number of species recorded around the islands reflect the diverse habitats they support and yet that also overlap with important fishing zones.

Only two species of deep-sea sharks were recorded in this study despite recent additions of seven new records from the region [14, 23–25]. This was due to the logistical difficulties in sampling the large quantities of deep-sea sharks landed, along with time constraints between landings and transport to the storage units. Currently, there is an ongoing targeted deep-sea shark fishery in the Andaman Islands that supplies the demand for shark liver oil [15]. Deep-sea sharks have rates of population increase, that are on average, less than half those of shelf and pelagic species and some of the lowest levels recorded to date [34]. These life history traits do not allow them to sustain intense fishing pressure which can lead to rapid population declines. This has been previously documented in the Indian Ocean region with the collapse of deep-sea fisheries along the west coast of India and the Maldives occurring within a short time period after the beginning of their exploitation [17, 35]. In addition, population recovery rates also decrease with increasing depth, suggesting that these species are most susceptible to overexploitation [34]. Thus, we emphasise the urgency and importance of assessing the status and monitoring the populations of deep-sea sharks as well as determining the socio-economic benefits and impacts of the trade in shark liver oil so that management measures such as catch limits, and spatial or temporal regulations can be put in place in order to avoid a collapse of this fishery.

Many rays (e.g., *Neotrygon* sp., *Pastinachus* sp.) could not be identified to the species level due to their tails being cut, difficulty in manipulation due to their weight, or traders transporting them before photo documentation was possible. Ongoing taxonomical uncertainty for many ray species currently exists in India, where there is ambiguity in several species complexes. In order to address and resolve this, a robust taxonomic framework needs to be developed which can be used to better understand diversity and potential impacts from fishing pressure on key species. Thus, in the future, a combination of molecular techniques, and long-term fishery-independent surveys need to be established to gain a holistic picture of diversity as well as population trends in the region.

At many sites sampled around the world, smaller size species are predominantly landed, as many of the larger shark species have been overfished [36–39]. On peninsular India, shark stocks have also declined over the past decade with smaller, faster-growing shark species displacing larger, slower-growing species [5, 11, 40 - 43]. A decrease in the diversity of species landed has also been documented in areas with high fishing pressure. Indeed, Thailand, closer to Andaman and Nicobar Islands than to mainland India, has recorded a decrease in landings of larger sharks from 41 species in 2004 to 15 species in 2014-15 [44]. Yet our results indicated that this is not yet the case in the Andaman and Nicobar Islands as four of the six dominantly landed sharks are larger bodied shark species. This suggests that we are still at a point where informed management decisions can lead to the conservation of these populations. However, as gravid females, juveniles and YOYs are being fished, the productivity, resilience and sustainability of these populations may have already reduced [45].

The largest size range in sharks was recorded in landings from pelagic longlines and gillnets. While gillnets fish up to seven nautical miles from the coast across the Andaman Islands, pelagic longlines fish exclusively beyond seven nautical miles from the coast and within 12 nautical miles, and are known to fish in waters from South Andaman to Nicobar. The high range of TL and non-specificity of gear catch could be ascribed to the gear size, fishing grounds, or the activity patterns of the diverse species ecology. In future, size - specificity studies in relation to the catch by gears need to be conducted in order to determine gear modifications best suited for the susceptible life history stages of threatened shark and ray populations.

This study emphasizes the overlap between critical habitats and fishing grounds as all life-stages for most species were recording highlighting their susceptibility to fishing pressure. Gravid females of 12 species were reported with fishers confirming that they were fished in the waters of the Andaman and Nicobar Islands. Juveniles of large shark species are being fished intensely, such as *Carcharhinus albimarginatus*, *C. amblyrhynchos*, *C. brevipinna* and *Sphyrna lewini*, which is a reason to be concerned as these species exhibit particularly low productivity and growth rates leading to high susceptibility to anthropogenic pressure and are slow to recover from overexploitation [46]. The presence of high abundance of YOY for these species suggests that these species might be using the islands as pupping or nursery grounds. *Carcharhinus brevipinna* and *S. lewini* have been recorded to use inshore nursery areas for their young [47–49]. Thus, we recommend that these breeding and nursery grounds need to be identified and evaluated, following which they can be temporally and spatially managed.

Sex ratios in landings differed across species and fishing gears, which could be due to confounding factors such as gear selectivity, fishing grounds, season, productivity, currents and bathymetry [51]. Significantly more females than males for *C. amblyrhynchos, S. lewini,* and *P. jenkinsii* suggests that females of these species dominate the populations in these waters. These are also aggregating species often exhibiting some degree of side fidelity [52–56] another ecological character that needs to be considered in spatial management. Similarly, for *L. macrorhinus,* and *H. leoparda,* significantly more males were landed than females, whereas parity was recorded for *C. falciformis*. In future, region-specific studies need to be carried out to assess sex-mediated spatial ecology for sharks and rays. Systematic sampling from fishing vessels across seasons would also be required to get fine-scale overlap between temporal and spatial distribution of shark and rays as well as fishing gear specificity.

Landings for sharks peaked from November to April, coinciding with pelagic longlines targeting sharks during this time. Landings in December were unpredictable where sampling differed from the highest number of sharks to a complete absence of sharks resulting in a higher standard error. Seasonal differences during the year could be ascribed to various factors such as the weather, access to fishing grounds, fishing gears used, and the ecology of the species fished. During the SW monsoon (May to September), the absence of landings at the Junglighat site could be due to the weather which makes it risky for fishers to go out fishing or the seasonal ban on trawlers and pelagic longliners.

It is noteworthy to highlight species diversity, quantities landed and TL ranges were highest in pelagic longlines. Landings from these gears included threatened species such as *Alopias pelagicus, A. superciliosus*, *C. falciformis*, *C. longimanus*, and *S. lewini* which are migratory species. These species are listed under Appendix I (*C. longimanus*) and II of the Convention on the Conservation of Migratory Species of Wild Animals (CMS) and the Convention on International Trade in Endangered Species of Wild Fauna and Flora (CITES), for international cooperation for conservation of migratory species and to regulate their trade, respectively. Since India is a signatory to these conventions, there is an urgent need for regional cooperation to ensure their protection as well as trade controls. CITES specifically requires the development of a Non-Detrimental Findings to assure that trade is not adversely impacting populations [59], something that has yet to be done in India. Given India’s long coastline of nearly 7,516 km, along with the multi-stakeholder and multi-gear nature of fisheries, it is challenging to comprehensively monitor the trends in landings of sharks and rays. While the Central Marine Fisheries Research Institute (CMFRI) in India has the most comprehensive fisheries database dating back to 1947, it is restricted to peninsular India, with no presence in the Andaman and Nicobar Islands. Here, the monitoring is undertaken by the Andaman and Nicobar Islands Directorate of Fisheries who broadly focuses on commercial fish stocks and does not include species-specific categories for sharks and rays [15]. Additionally, the Zoological Society of India (ZSI), Fisheries Survey of India (FSI) and Central Island Agricultural Research Institute (ICAR) conduct opportunistic surveys to document species diversity. We conducted this study in the Andaman Islands to fill this gap, however, additional studies are required to address ongoing taxonomic ambiguities, improve knowledge of species by expanding fisheries independent monitoring, and to facilitate long-term species-specific monitoring. The latter would benefit the government as it would ensure traceability and control of onward trade. This in turn could help determine management and conservation measures for implementing CITES.

Shark and ray species protected under the WLPA were rarely landed (only two individuals of *U. asperrimus* were recorded). Most of the species listed in the WLPA are found in estuarine habitats and are not likely to occur around the islands, including *Anoxypristis cuspidata*, *Glyphis gangeticus*, and *G. glyphis*. *Rhynchobatus djiddensis* listed in the WLPA does not appear to occur in India and the species complex could include *R. australiae* and *R. laevis* [60]. However, the latter two species are not protected under the WLPA. Anecdotal reports from fishers state that a few of these species (e.g., *Pristis* sp.) have not been seen or landed for over a decade (Z. Tyabji unpubl. data). This highlights the urgent need for amending the WLPA and to include Critically Endangered and Endangered species that occur in India to the list of protected species. However, species-selective bans in non-selective multi-gear fishery are difficult to implement, thus amending the WLPA has to be combined with stakeholder engagement and other regulations such as fishing gear modifications and spatial closures.

While there exists a 45-day shark fishing ban, there are no regulations for ray fishing, despite them being predominantly threatened species. Of the 19 ray species identified, 15 species (85.17 %) are listed on the IUCN Red List of Threatened Species as threatened (Critically Endangered, Endangered, or Vulnerable), one species (0.4 %) as Near Threatened, one species (1.08 %) as Data Deficient, and two species (13.33 %) have not been evaluated. Rays are extremely susceptible to overexploitation, with wedgefishes and giant guitarfishes being the most imperiled marine taxa globally [1, 60]. Susceptibility studies on the various sharks and ray species in Papua New Guinea, deemed *P. jenkinsii* at the highest risk in trawl fisheries [61]. This was one of the most dominant species landed in the Andaman and Nicobar Islands. This is concerning as most ray species utilize coastal areas which overlap with the majority of fisheries. Additionally, there is a developing targeted ray fishery in the islands (Z. Tyabji unpubl. data) due to the local demand for their meat and trade in their skins. Studies regarding the local population status and exploitation rate of rays on the islands are urgently required, following which a prioritizing exercise needs to be conducted which takes into account the life history traits, susceptibility to fishing pressures, and population recovery rate. Based on this, ray species that are most susceptible to overexploitation need to be identified and a management plan needs to be developed and implemented.

While sustainability can be attained by a combination of policy changes such as the identification and protection of critical shark and ray habitats and populations, gear modifications, and implementing seasonal and temporal bans, it is a daunting task due to the lack of data on which to base these management strategies. We recommend additional studies and continued long-term monitoring with a focus on threatened species in order to establish appropriate management measures. We also need to understand the socio-economic importance of shark and ray fisheries for the range of stakeholders and communities on the islands; and the role of these fisheries in the supply chain of both domestic and global markets while designing management strategies. It is essential that policy formulation and changes are carried out with the involvement of fishers and local stakeholders for effective implementation. Thus, we suggest adapting science-based management techniques with the inclusiveness of stakeholders involved so as to avoid overexploitation of sharks and rays and aid in their conservation.

## ACKNOWLEDGEMENT

We would like to thank the fishers, traders and processing unit managers at the study sites for their help and support. We are grateful to Nairika Barucha, Vishwanath K. G., Harsh Narola, Mahi Mankeshwar, Anushka Rege, Evan Nazareth, Mahadev, Sachin Vaishampayan, and Sitara Hussain for helping with data collection. Lastly, we are thankful to Andaman Nicobar Environment Team for their logistical support.

## SUPPORTING INFORMATION

**S1 Table. Maximum likelihood estimates of length and weight regression parameters for the six commonly landed shark species.**

**S1 Fig. 1) *Aetobatus flagellum*** (a) dorsal view (b) ventral view of the mouth; **2) Two colourations of *Himantura tutul*** (a) dorsal view (b) denticles on the nuchal area (c) dorsal view (d) denticles on the nuchal area; **3) *Pateobatis fai*** (a) dorsal view (b) ventral view (c) tail

**S2 Fig. Sharks and rays landed at the fish landing sites.** Clockwise from top left: Deep-sea sharks caught from deep-sea longline landed at Burmanallah; Fishers take out sharks from the pelagic longline boats at Junglighat; Shark fins kept to dry; Landed rays are weighed, following which they will be transported to the storage units; Adult and juvenile sharks of various species landed at Junglighat.

## AUTHOR CONTRIBUTIONS

Conceptualization: ZT, VP, DS

Data Curation: ZT

Formal Analysis: ZT

Funding Acquisition: ZT

Investigation: ZT, TW

Methodology: ZT, RWJ, DS

Project Administration: ZT

Resources: ZT

Software: ZT, RWJ

Supervision: RWJ, DS

Validation: RWJ, DS

Visualization: ZT, TW

Writing – Original Draft Preparation: ZT

Writing – Review & Editing: ZT, TW, VP, RWJ, DS

